# Age-induced midbrain-striatum assembloids model early phenotypes of Parkinson’s disease

**DOI:** 10.1101/2023.10.28.564305

**Authors:** Kyriaki Barmpa, Claudia Saraiva, Gemma Gomez-Giro, Elisa Gabassi, Sarah Spitz, Konstanze Brandauer, Juan E. Rodriguez Gatica, Paul Antony, Graham Robertson, Florentia Papastefanaki, Ulrich Kubitscheck, Ahmad Salti, Peter Ertl, Rebecca Matsas, Frank Edenhofer, Jens C. Schwamborn

**Author notes:** Corresponding author: Jens C. Schwamborn.

## Abstract

Parkinson’s disease (PD), one of the most common aging-associated neurodegenerative disorders, is characterised by nigrostriatal pathway dysfunction, caused by the gradual loss of dopaminergic neurons in the substantia nigra pars compacta (SNpc) of the midbrain and the dopamine depletion in the striatum. State of the art, human *in vitro* models are enabling the study of the dopaminergic neurons’ loss, but not the dysregulation of the dopaminergic network in the nigrostriatal pathway. Additionally, these models do not incorporate aging characteristics which potentially contribute to the development of PD. Therefore, it is conceivable that research conducted using these models overlooked numerous processes that contribute to disease’s phenotypes. Here we present a nigrostriatal pathway model based on midbrain-striatum assembloids with inducible aging. We show that these assembloids are capable of developing characteristics of the nigrostriatal connectivity, with catecholamine release from the midbrain to striatum and synapse formation between midbrain and striatal neurons. Moreover, Progerin-overexpressing assembloids acquire aging traits that lead to early phenotypes of PD. This new model shall help to reveal the contribution of aging as well as nigrostriatal connectivity to the onset and progression of PD.

## Introduction

The nigrostriatal pathway connectivity is established through projections of the midbrain SNpc dopaminergic neurons to the dorsal striatum. Disruption of the nigrostriatal connectivity contributes to the development of PD symptoms, such as motor dysfunction, tremor, muscle stiffness, and bradykinesia (Caminiti et al., 2017; Ghosh et al., 2019). The gradual loss of dopaminergic neurons leads to dopamine depletion in the striatum, with consequences in the synaptic network organization and cellular complexity (Zhai et al., 2018). Before the onset of neuronal loss there is significant decrease of dopaminergic axons in the striatum. This reduction in dopaminergic axons is twice as pronounced in the striatum compared to the decrease in dopaminergic neuronal cells in the SNpc (Chung et al., 2020; Heng et al., 2023; L. H. Li et al., 2009; Tagliaferro & Burke, 2016). However, due to the research focus on the loss of neurons, which is the ultimate endpoint of the disease, the mechanisms of the retrograde dopaminergic axons’ degeneration from the striatum are still not understood (Tagliaferro & Burke, 2016).

Animal models of PD have been used to elucidate cellular and molecular dysfunctions, but they cannot fully recapitulate the impairments that occur in the human nigrostriatal connectivity (Potashkin et al., 2011). More recently, human based cellular models such as induced pluripotent stem cells (iPSCs) derived 2D neuronal cell culture and the more advanced 3D midbrain organoids have been extensively used for PD modelling. Midbrain organoids have the advantage of better recapitulating the neuronal connectivity in a 3D environment with active neuronal electrophysiology and self-organization properties (J. Kim et al., 2020; Monzel et al., 2017). Midbrain organoids derived from PD patient cell lines exhibit phenotypes relevant to the disease and they present a great tool for further understanding the molecular and cellular mechanisms of neurodegeneration (Becerra-Calixto et al., 2023; Jarazo et al., 2019; Jo et al., 2021; H. Kim et al., 2019; S. W. Kim et al., 2021; Smits et al., 2019; Zagare et al., 2022). Although these models are useful for studying the dopaminergic neurons’ vulnerability in PD, they do not recapitulate the nigrostriatal pathway connectivity dysfunction which is essential for unveiling the sources of PD onset and progression.

More advanced 3D models with the combination of two or more region specific brain organoids have started to pave the way towards cellular models with higher complexity that can effectively reflect the neuronal connectivity between different brain regions (Makrygianni & Chrousos, 2021; Panoutsopoulos, 2020; Pașca, 2018). For instance, recent studies have shown the generation of pallium-subpallium, cortico-motor and cortico-striatal assembloids from human iPSCs (Andersen et al., 2020; Birey et al., 2017; Miura et al., 2020), and their capability to develop physiological functional networks that could provide essential insights in diseases’ development and progression.

Here we present the development of a midbrain-striatum assembloid model that can recapitulate the nigrostriatal pathway connectivity, with catecholamines release from the midbrain to striatum and the formation of active synapses between the midbrain dopaminergic neurons and the striatal neurons. Although the midbrain-striatum assembloid model allows us to study aspects of the human nigrostriatal connectivity *in vitro*, it still lacks one major risk factor of PD, which is aging (Collier et al., 2017; Hindle, 2010; Lapasset et al., 2011). To address whether aging induction could lead to neurodegenerative phenotypes in assembloids, we used a genetically engineered iPSC line that carries a Progerin-GFP transgene under the control of the Tet-On system for the controllable overexpression of Progerin in the presence of doxycycline (Gabassi et al., *In Preparation*). Our results show that Progerin-overexpression can induce aging phenotypes in the midbrain-striatum assembloid model which subsequently leads to the development of PD-associated early neurodegeneration phenotypes.

## Methods

### iPSCs and NESCs

The induced pluripotent stem cells (iPSCs) that were used in this study are described in Supplementary Table 1. The iPSCs were cultured in 6-well plates (Thermo Fisher Scientific, 140675) coated with Geltrex (Life Technologies, A1413302). For the first 24 hours the cells were cultured in Essential 8 Basal medium (Thermo Fisher Scientific, A1517001) supplemented with 1% Penicillin/Streptomycin (Invitrogen, 15140122) and 10 µM ROCK Inhibitor (Ri) (Y-27632, Millipore, SCM075). After the 24 hours, the cells were cultured in Essential 8 Basal Medium, with daily media changes. Confluent iPSCs (∼70-90%) were splitted using UltraPure™ 0.5M EDTA, pH 8.0 (Thermo Fisher Scientific, 15575020). Immunofluorescence staining was performed to confirm the pluripotent identity of the iPSCs (Supplementary Figure 1.A-D). Neuroepithelial stem cells (NESCs) were generated from iPSCs using a previously described protocol (Reinhardt et al., 2013). Similar to the iPSCs, the NESCs were cultured in Geltrex-coated 6-well plates using supplemented N2B27 media, as described in (Monzel et al., 2017). The NESCs were passaged using Accutase (Sigma-Aldrich, A6964) and their neural stem cell identity was confirmed with immunofluorescence staining (Supplementary Figure 1.E-G).

### Striatum Organoids

Striatum organoids (StrOs) were generated using an adapted protocol (C4) from (Miura et al., 2020) (C3) after comparing it with two other conditions (RA and SR) in two time points of culture (day (D)35 and D50) (Supplementary Figure 2, Supplementary Figure 3). iPSCs at ∼70% confluency were used for the StrOs generation. Before the procedure of spheroids formation, the cells were treated overnight with 1% DMSO (Sigma-Aldrich, D2650) in Essential 8 basal medium. For the spheroids formation (D-1), the iPSCs were first incubated into Accutase at 37 °C for 5 min. The accutase was stopped using 5x DMEM-F12 (Thermo Fisher Scientific, 21331-046). The cells were then resuspended in Essential 8 medium containing 20 μM Ri and counted using the Countless cell counting chambers slides (Invitrogen, C10313). For condition C4, 10000 cells were added per well in the BIOFLOAT™ 96-well plate U-bottom (faCellitate, F202003), centrifuged in 100g for 3 min and then incubated in the normal culture conditions of 37°C with 5% CO2. The spheroids were left intact for two days to form properly and at D1 the medium was exchanged with Essential 6 medium (Thermo Fisher Scientific, A1516401) supplemented with 10 μM Ri, 2.5 μM Dorsomorphin (Sigma-Aldrich, P5499) and 10 μM SB-431542 (Abcam, ab120163). The spheroids were cultured in the same medium until D5, with a reduction of the Ri concentration at 10 μΜ at D2, 5 μΜ at D3 until completely removed at D4. To start the differentiation process of the spheroids into StrOs, on D6 the medium was exchanged with a medium containing Neurobasal-A (Thermo Fisher Scientific, 10888022), 2% B-27 without vitamin A (Thermo Fisher Scientific, 12587010), 1% Penicillin/Streptomycin (Invitrogen, 15140122), 1% GlutaMAX (Thermo Fisher Scientific, 35050061) and supplemented with 2.5 μM IWP-2 (Selleckchem, S7085) and 50 ng/ml Activin A (Thermo Fisher Scientific, PHC9561). From D9 to 17, the media was additionally supplemented with 100 nM SR11237 (Tocris, 3411). Until D17 the medium was exchanged daily. From D17 to 35, the media was changed to promote the neuronal differentiation and it was supplemented with 20 ng/ml BDNF (PeproTech, 450-02), 20 ng/ml NT-3 (Alomone labs, N-260), 200 μM AA (Sigma-Aldrich, A4544) and 100 μM cAMP (Biosynth, ND07996), with medium exchanges every 3 to 4 days. For condition C3, the same protocol as described in Miura and colleagues (Miura et al., 2020) was followed, without the addition of cis-4,7,10,13,16,1 9-docosahexaenoic acid (DHA) at D22 of differentiation.

For the generation of StrOs from the RA and SR conditions, iPSCs at ∼70% confluency were collected using accutase as described above, and 9000 cells were plated in each well of the BIOFLOAT™ 96-well plate U-bottom (D-2). The media used for plating was containing 80% DMEM F12 (Thermo Fisher Scientific, 21331046), 20% KOSR (Thermo Fisher Scientific, 10828028), 3% FBS (Invitrogen, 16140071), 1% GlutaMAX, 1% NEAA (Thermo Fisher Scientific, 11140-050) and 0.7% 2-Mercaptoethanol 50 mM (Thermo Fisher Scientific, 31350-010). This media was supplemented with 10 μM Ri and 40 ng/ml FGF-basic (PeproTech, 100-18B). After two days (D0 of culture) the media was exchanged with the Neural induction medium (NIM) containing 76.8% DMEM F12, 20% KOSR, 1% NEAA, 1% GlutaMAX, 1% Penicillin/Streptomycin and 0.2% 2-Mercaptoethanol 50 mM. From D0 to 2 the NIM was supplemented with 5 μΜ DM, 10 μM SB and 10 μM Ri, while on D2 Ri was removed. From D2 to D16 the medium was exchanged daily. On D4 and 5 the medium was additionally supplemented with 5 μΜ IWP-2. On D6, NIM was exchanged to the neural differentiation medium (NM) which contained 96% Neurobasal A medium, 2% B-27 without vitamin A, 1% GlutaMAX and 1% Penicillin/Streptomycin. From D6 to 8, NM was supplemented with 20 ng/ml FGF-basic, 20 ng/ml EGF (PeproTech, AF-100-15) and 5 μΜ IWP-2. From D8 to 16, NM was supplemented with 20 ng/ml FGF-basic, 20 ng/ml EGF, 5 μΜ IWP-2, 50 nM SAG (Merck Millipore, 566660), 50 ng/ml Activin A and 1 mM for condition RA or 100 nM SR11237 for condition SR. On D17 the media was exchanged with a non-supplemented NM. From D18 to D35 or D50, NM was supplemented with 20 ng/ml BDNF and 20 ng/ml NT3, with media changes every 3 to 4 days.

### Midbrain organoids

Midbrain organoids (MOs) were generated using NESCs as the starting population of cells. The protocol that was used is a slightly altered version of the one described in Monzel and colleagues (Monzel et al., 2017). At D0, 9000 NESCs were added per well in the BIOFLOAT™ 96-well plate U-bottom and cultured in maintenance medium for 2 days. At D3 the medium was exchanged with the differentiation medium containing 1 µM purmorphamine and at D8 with the final differentiation medium without purmorphamine. The organoids were kept in static conditions and in the 96-well plates U-bottom, until used for the assembloids generation (described below).

### Assembloids

Assembloids of midbrain and striatum organoids were generated using midbrain organoids at D20 and striatum organoids at D35 of culture. Due to the different media composition of the organoids, an optimization of the co-culture medium was needed (Supplementary Figure 4, Supplementary Figure 5, Supplementary Figure 6). Four different media were tested. The Neural Medium (NM) was comprised of Neurobasal-A, 2% B-27 without vitamin A, 1% Penicillin/Streptomycin and 1% GlutaMAX, while the Neural Medium Plus (NMpl) was supplemented with B-27 plus (Thermo Fisher Scientific, A3582801) instead of the B-27 without Vitamin A. In the third condition, the Neural Medium++ (NM++) was tested, which was the NM medium supplemented with 20 ng/ml BDNF (PeproTech, 450-02), 10 ng/ml GDNF (PeproTech, 450-10), 20 ng/ml NT-3 (Alomone labs, N-260), 200 μM AA (Sigma-Aldrich, A4544) and 100 μM cAMP (Biosynth, ND07996). The optimal assembloid co-culture condition for the assembloid model was consisted of the N2B27 medium (described by Monzel and colleagues (Monzel et al., 2017)), which consists of DMEM F12 (Invitrogen)/Neurobasal (Invitrogen) 50:50 with 0.5% N2 supplement (Thermo Fisher Scientific, 17502001), 1% B-27 without Vitamin A, 1 % GlutaMAX and 1 % Penicillin/Streptomycin. The media was further supplemented with 20 ng/ml BDNF (PeproTech, 450-02), 10 ng/ml GDNF (PeproTech, 450-10), 20 ng/ml NT-3 (Alomone labs, N-260), 200 μM AA (Sigma-Aldrich, A4544) and 100 μM cAMP (Biosynth, ND07996). At D0 of the assembloids generation, each StrO was transferred in each well of the 96-well plate U-bottom that contained the MOs and the medium was exchanged to the assembloid co-culture medium. After 4 days, the two organoids were merged into an assembloid, and they were transferred in 24 well ultra-low attachment plates (Celltreat, 229524). Some of the assembloids were embedded in 30 µl Geltrex (Invitrogen, A1413302), as described before (Monzel et al., 2017). The assembloids were cultured in 37°C, 5% CO2 under static conditions.

For the Progerin overexpression induction, assembloids that were generated from the Progerin-overexpressing cell line were treated with 4 ng/μl doxycycline (Sigma-Aldrich, D9891). Generation and validation of iPSC line expressing Progerin under control of the Tet-ON system is described elsewhere (Gabassi et al., *In Preparation*).

### ATP and LDH assay

Intracellular ATP in assembloids was measured using luminescence based CellTiter-Glo® 3D Cell Viability Assay (Promega, G9681). Three organoids per cell line and per condition were transferred each in one well of the imaging plate (PerkinElmer, 6055300). 50 µl of CellTiter-Glo® reagent were added to each well and the plate was incubated for 30 min on a shaker at room temperature (RT). Luminescence was measured using Cytation5 M cell imaging reader (RRID:SCR_019732). The experiment was repeated for three independent derivations (batches) at D30 assembloids. The mean signal of three assembloids of each cell line and condition was calculated and normalized to the mean size (area) of the assembloids. Brightfield images of random assembloids of the three batches taken with the ZEISS Axio Vert.A1+Axiocam ICM1 microscope and the area of each assembloid was calculated with the ZEN (blue edition) software (RRID:SCR_013672). The mean area of all assembloids measured per line and conditions from each batch was used for the normalization.

For determining the cytotoxicity in the assembloids, the LDH-Glo™ Cytotoxicity Assay (Promega, J2381) was used. In the day of assembloids collections (D30 of culture) media from three assembloids per line and condition from three batches was collected and snap frozen in liquid nitrogen. For the LDH assay, the snap frozen media was thawed on ice. 50 μl of media from each sample and 50 μl of the enzyme and substrate mix was pipetted in a well of the imaging plate (PerkinElmer, 6055300). The plate was briefly mixed and incubated for 1 h at RT avoiding exposure to light. Luminescence was measured using Cytation5 M cell imaging reader. Similar to the ATP assay, the mean signal of the assembloids was normalized to the mean of the assembloids’ area.

### Western Blotting – RIPA buffer

For Western blotting, four non-embedded assembloids or six to eight organoids were lysed using RIPA buffer (Abcam, ab156034) supplemented with cOmplete^TM^ Protease Inhibitor Cocktail (Roche, 11697498001) and Phosphatase Inhibitor Cocktail Set V (Merck Millipore, 524629). The samples were pipetted 10-20 times up and down until dissolved and were incubated on ice for 20 min. For DNA disruption, lysates were sonicated for 10 cycles (30 seconds on / 30 seconds off) using the Bioruptor Pico (Diagenode), followed by a centrifugation at 4°C for 20 min at 14000g. The protein concentration was measured using the Pierce™ BCA Protein Assay Kit (Thermo Fisher Scientific, 23225). Samples were adjusted to the same concentration by appropriate dilution with RIPA buffer and boiled at 95°C for 5 min in denaturating loading buffer. 2.5-10 μg of protein was loaded per sample for every Western blot. Protein separation was achieved using SDS polyacrylamide gel electrophoresis (Bolt™ 4-12% Bis-Tris Plus Gel, Thermo Fisher Scientific) and transferred onto a PVDF membrane using iBlot™ 2 Gel Transfer Device (Thermo Fisher Scientific). After transfer the membrane was dried for 10 min in 37°C and subsequently activated with 100% Methanol for 30 sec. The membranes were washed twice with PBS containing 0.02% Tween and they were blocked for 1 hour at RT in 5% skimmed milk powder dissolved in PBS. After blocking, the membranes were washed quickly with PBS containing 0.02% Tween and were incubated overnight at 4°C with the primary antibodies prepared in 5% BSA and 0.02% Tween in PBS (Supplementary Table 2). The next day, membranes were washed three times for 5 min with PBS containing 0.02% Tween and incubated with DyLight™ secondary antibodies at a dilution of 1:10000 (anti-rabbit IgG (H+L) 800, Cell Signaling, 5151P or anti-mouse IgG (H+L) 680, Cell Signaling, 5470P) for 1 hour. Membranes were revealed in the Odyssey® Fc 2800 Imaging System and exposure time was from 30 sec to 4 min, depending on the primary antibody used. Western blots were analyzed using ImageJ (RRID:SCR_003070) software.

### Western Blotting – Nuclear and Cytoplasmic fractionation

The protocol described by Abcam (https://www.abcam.com/protocols/nuclear-extraction-protocol-nuclear-fractionation-protocol) was used for the nuclear-cytoplasmic fractionation. At the last step of the protocol both nuclear and cytoplasmic samples were sonicated for 10 cycles (30 seconds on / 30 seconds off) using the Bioruptor Pico (Diagenode). For the preparation of the nuclear extraction and fractionation buffer the reagents used were HEPES (Sigma-Aldrich, H3375), KCl (AppliChem, 8059), MgCl2 (Sigma-Aldrich, M8266), EDTA (Sigma-Aldrich, E9884), EGTA (Sigma-Aldrich, E3889), DTT (Thermo Fisher Scientific, R0861), cOmplete^TM^ Protease Inhibitor Cocktail and Phosphatase Inhibitor Cocktail Set V. The following Western blotting procedure is described in the previous section.

### Flow cytometry

Three embedded assembloids per condition were used for GFP+, live cells measurement in BD LSRFortessa flow cytometer (RRID:SCR_019601). Geltrex embedded assembloids were first incubated at 37°C for 40-50 min on shaker in 500 μl of papain solution containing 20 ml DMEM-F12, 36 mg Papain (Sigma-Aldrich, P4762), 8 mg EDTA (Sigma-Aldrich, E6758) and 8 mg L-Cystein (Sigma-Aldrich, C6852). To start the dissociation process, papain solution was replaced with 500 μl accutase and the assembloids were pipetted with the 1000 pipette, followed by a 10 min incubation shaking. After that, pipetting with the 200 μl pipette and incubation cycles were continued until the complete dissociation of the assembloids. For the accutase and papain inhibition, 500 μl papain inhibitor solution containing 5 mg/ml BSA (Carl Roth, 8076.4) and 5 mg/ml Trypsin inhibitor (Sigma-Aldrich/Roche, 10109878001) in PBS was added. After transferring the total volume in 2 ml Eppendorf tube, the dissociated assembloids were centrifuged at 500xg for 5 min. Supernatant was discarded and the pellet was washed once with PBS. The pellet was resuspended in 300 μl DMEM (Thermo Fisher Scientific, A14430-01) containing 1:1000 concentration live-dead stain Zombie NIR (Biolegend, 423106), followed by incubation at 37°C for up to 30 min. Cells were then centrifuged at 500xg for 3 min and pellet was washed twice with PBS and centrifuged again with the same setting. After the final wash and centrifugation, the pellet was resuspended in DMEM and the samples were run in Becton Dickinson LSRFortessa, with 10000 events acquisition of GFP+, live-cells. Each sample was run in two technical replicates. The data were analysed using the FlowJo software (v.10.7.2, RRID:SCR_008520).

### Rabies virus based retrograde monosynaptic tracing

#### Lentiviral vector and rabies virus vector productions

The construct pBOB-synP-HTB [gift from Edward Callaway & Liqun Luo (Addgene plasmid # 30195; http://n2t.net/addgene:30195; RRID:Addgene_30195)] (Miyamichi et al., 2011) was used for the production of the replication-deficient LV-GP-TVA-GFP lentiviral vector. High-titer preparations of lentiviral particles were produced, as previously described (Kutner et al., 2009). The titer of the preparation used was 3 × 10^8^ IU/mL.

For the production of RBV-ΔG-EnvA-RFP rabies viral particles, we followed stages III-VI of the previously established protocol for the amplification, pseudotyping, and concentration of the virus (Osakada & Callaway, 2013). Titer was 4 × 10^7^ IU/mL.

#### Lentiviral vector and rabies virus vector transduction of assembloids

To assess the connectivity through active synapses between the midbrain and striatum neurons in the assembloid model, striatum organoids at D35 of culture were transduced by adding concentrated LV-GP-TVA-GFP viral particles at 1:500 dilution, in the culture medium. After 7 days the medium containing the lentiviral vector was discarded and the organoids were washed twice with fresh medium. The LV-transduced StrOs were then merged with MOs and the assembloids were infected by the addition of the concentrated RBV-ΔG-EnvA-RFP rabies viral particles at 1:500 dilution in the culture medium. Control experiments, with single infection of assembloids with only one of the vectors at a time, were performed to confirm the specificity of the signal. After 7 days the media was changed and the assembloids were cultured for up to D30. 70 μm-thick sections were either imaged directly for the observation of the RFP and GFP signal from the viral infections or were immunostained using a tyrosine hydroxylase (TH) antibody (see **Immunofluorescence Staining** procedure and Supplementary Table 3 for the antibody) to assess the colocalization between TH and RFP.

### β-galactosidase

70 μm sections from assembloids were used in the β-galactosidase staining, using the Senescence Detection Kit (Abcam, ab65351). One section from two or three assembloids per condition from four batches were used. Images were taken using the colour camera setting with 4X objectives in the Olympus IX83 microscope (RRID:SCR_020344). β-galactosidase positive areas were quantified using ImageJ. Positive areas in each section were summed and normalised to the total area of the section in the image. The normalised value was multiplied by 100 for calculating the % of positive β-galactosidase areas in the image.

### Immunofluorescence Staining iPSCs and NESCs

The procedure that was used for the immunofluorescence staining characterization of the iPSCs is detailed described by Gomez-Giro and colleagues (Gomez-Giro et al., 2019), a previous study from our lab.

For the NESCs’ immunofluorescent staining characterization, NESCs were cultured on Geltrex-coated 96-well imaging plates (PerkinElmer, 6055300) until they reached approximately 70% confluency. Next, NESCs were fixed for 15 min at RT with 4% Paraformaldehyde (PFA), washed 3x for 5 min with PBS and permeabilizated with 0.3% Triton X-100 in PBS for 15 min at RT. After permeabilization, NESCs were washed 3x for 5 min with PBS and then blocked with 10% fetal bovine serum (FBS) in PBS for 1 hour at RT. Primary antibodies (Supplementary Table 3) were diluted in 3% FBS in PBS and the cells were incubated overnight at 4°C. Then cells were washed 3x for 5 min with PBS and incubated for 1 hour at RT with secondary antibodies (Supplementary Table 3) and Hoechst 33342 (Invitrogen, 62249). After 3 washes for 5 min with PBS, cells were kept in 0.1% Sodium Azide (NaAz) in PBS until imaging with confocal microscopy.

### Organoid and Assembloid Sections

Assembloids and organoids were fixed with 4% PFA overnight at 4°C, and then washed with PBS three times for 15 min at RT. At least three organoids/assembloids per line and time point were embedded in 3% low-melting point agarose (Biozym, 840100). 70 μm sections were obtained using the vibratome (Leica VT1000s, RRID:SCR_016495). The sections were permeabilized for 30 min in 0.5 % Triton X-100 and blocked for 2 h with blocking buffer containing 2.5% normal goat serum, 2.5 % BSA, 0.01% Triton X-100 and 0.1 % sodium azide in PBS at RT. Sections were incubated with the primary antibodies (Supplementary Table 3) diluted in blocking buffer for 48-72 hours at 4 °C. The sections were washed with 0.01% Triton X-100 for 5 min three times and then incubated with the secondary antibodies (Supplementary Table 3) and Hoechst at 1:1000 dilution for 2 hours at RT. The sections when then washed again with 0.01% Triton X-100 for 5 min three times at RT. After the last wash, the sections were kept in MilliQ water and mounted on slides as described by Nickels and colleagues (Nickels et al., 2020).

### Microscopy

For high-content image analysis, one 70 μm section from three organoids/assembloids of each condition from at least three batches were acquired using the Yokogawa CV8000 high content screening microscope (RRID:SCR_023270) with a 20X/0.75 numerical aperture (NA) objective.

For qualitative analysis, images were acquired using a confocal laser scanning microscope (Zeiss LSM 710, RRID:SCR_018063) with the 20X/0.8 NA, 40X/1.3 NA or 63X/1.4 NA objective.

### Light sheet fluorescence expansion microscopy

#### Sample preparation

The expansion microscopy protocol was adopted from Rodriguez-Gatica and colleagues (Rodriguez-Gatica et al., 2022) and used for whole assembloid preparation. Briefly, the labelled samples were incubated with 2 mM of methylacrylic acid-NHS linker for 24 hours on a shaker at RT, ensuring they were fully submerged in 1 ml of the solution. Following this, the samples were washed thrice, each time for 40 min in PBS. Subsequently, they were incubated in the monomer solution (comprising 8.6% sodium acrylate, 2.5% acrylamide, 0.15% N,N’-methylenebisacrylamide, and 11.7% NaCl in 1× PBS) on a shaker at 4°C, using 0.9 ml of the solution to cover them.

The gelling solution was freshly prepared by adding 4-hydroxy-TEMPO (0.01%), TEMED (0.2%), and ammonium persulfate (0.2%) to new monomer solution. To prevent premature polymerization during the gelling process, the samples were placed in a 24-well plate on ice. After discarding the monomer and adding the gelling solution, the samples were shaken at 4°C for 5 min. They were then transferred to a gelling chamber that utilized 2 mm spacers to maintain the sample’s integrity and prevent deformation. Post-transfer, the samples were left at 4°C overnight and subsequently incubated at 37°C for 2 hours. Upon gel formation, the samples were extracted from the gelling chamber and incubated at 37°C in the digestion buffer (containing 50 mM Tris, 1 mM EDTA, 0.5% Triton X-100, 0.8 M guanidine HCl, and 16 U∕ml of proteinase K; pH 8.0). The digestion buffer was replaced every 24 hours, if required, until the samples became fully transparent. After digestion, the buffer was discarded, and the samples were washed three times with PBS. The samples were then stored in PBS, resulting in an expansion to approximately 1.5 times their original size.

The ability to preserve the fluorescence of autofluorescent proteins during the expansion procedure allowed for exclusive immunolabeling of the dopaminergic neurons. Owing to the enhanced retention capabilities of the protocol, nuclear staining was conducted post-digestion. To ensure complete penetration throughout the gel, samples were incubated with Hoechst 33342 (H3570, Invitrogen) in PBS for 24 hours at RT with a concentration of 2.5 μg/ml (1:4000).

#### Light sheet microscopy

Following the expansion protocol, whole-assembloids were imaged using light sheet microscopy to obtain information at both the mesoscale and microscopic scale. Specifically, we utilized the Blaze microscope LaVision-Miltenyi Biotec for whole-assembloid imaging and a custom-built setup for higher resolution imaging of selected regions of interest (ROIs).

#### Mesoscopic Imaging

Mesoscopic imaging and analysis provide insights into the topology of complete assembloids. This is particularly useful for analysing the distribution of dopaminergic neurons throughout the entire sample. For imaging, the digested specimen was affixed to a coverslip using poly-L-lysine to prevent movement during measurement. This coverslip was then inserted into a custom sample holder and positioned within a large imaging chamber (183 × 50 × 64 mm length–width–height) filled with PBS solution (1xPBS, 0.02% Sodium azide).

For mesoscopic imaging, the Blaze from LaVision-Miltenyi BioTec was employed. Fluorescence excitation was achieved using four fiber-coupled lasers emitting at 405, 488, 561, and 638 nm (from the LaVision laser beam combiner). Given the sample’s transparency, a single illumination arm sufficed. The beam waist in the object plane was adjusted to a 1/e² diameter of 6 µm for all laser lines. Fluorescence emission was detected by a 4X objective (LaVision-Miltenyi BioTec MI PLAN 4X/0.35 NA, working distance (WD) 15 mm with water dipping cap) and data was captured using a sCMOS camera (2048 x 2048 pixels, pixel size 6.5 μm) in global shutter mode, resulting in an effective field of view of (3328 μm)². Due to the sample’s extensive size, mosaic-style imaging was essential to capture the entire organoid.

#### Microscopic Imaging

For microscopic scale imaging, assembloids were prepared as described above, producing a transparent specimen expanded by 1.5-fold. These were then examined using a high-resolution, long-distance objective with an NA of 1.1 (Nikon CFI75 25x/1.1 NA WD 2mm water immersion (WI)), achieving an optical resolution of approximately 0.3 µm laterally and 1.1 µm axially. Factoring in the sample expansion, this translated to an effective resolution of 0.2 and 0.7 µm, respectively, enabling structural characterization at subcellular length scales. For samples where the projections were not clear with the 4X mesoscopic view, a 12X objective (LaVision-Miltenyi BioTec MI PLAN 4X/0.35 NA, WD 8.5 mm WI) was used as an intermediate imaging step with the Ultramicroscope.

Imaging of complete assembloids at this resolution is often not advisable due to the data generation (about 500 Gigabyte data per 1 mm^3^ per channel). Instead, specific ROIs identified in the mesoscale data were chosen for subsequent microscopic scale analysis. To this end we employed a custom-built setup (Rodriguez-Gatica et al., 2022). Briefly, fluorescence excitation was achieved using four fiber-coupled lasers emitting at 405, 488, 561, and 638 nm (Hübner Photonics, Germany). A horizontally scanned light sheet was produced by a galvanometer system with silver-coated mirrors. Beam waist adjustment within the sample chamber was facilitated by relay optics mounted on a precision linear stage. The beam waist in the object plane was set to specific 1/e² diameters for each laser line. For illumination, a Mitutoyo 10x NA 0.28 air objective was used.

Our custom sample chamber (165 × 60 × 40 mm length–width–height) featured an illumination window created by a standard 24×24 mm coverslip with a 0.17 mm thickness. Observation was from above using a Nikon 25X 1.1 NA WI objective with an additional 1.5x magnification (Nikon, Germany). The sample, mounted on a coverslip, was manoeuvrable in three spatial directions via motorized micro-translation stages. Data capture was facilitated by a sCMOS camera (3200 x 3200 pixels, pixel size 6.5 μm, Kinetix, Teledyne Photometrics, USA), in global shutter mode, resulting in an effective field of view of (832 μm)². A custom-written LabView program managed all electronic components.

#### Data processing

3D stacks of raw 16-bit images were processed using custom-written MATLAB scripts, facilitating parallel data processing. Selected image stacks were spatially deconvolved using Huygens (Professional version 22.04, Scientific Volume Imaging, The Netherlands). This deconvolution utilized theoretical point spread functions (PSFs) derived from the microscopy parameters. The classical maximum likelihood estimation algorithm was employed, with Acuity set to: −10 to 25. Signal-to-noise ratio (SNR) values ranged between 12 and 20, and the maximum number of iterations varied from 60 to 100.

Full 3D representations of the samples were achieved by stitching multiple 3D datasets together using FIJI (Schindelin et al., 2012) and the stitching plugin by Preibisch and colleagues (Preibisch et al., 2009). To optimize the stitching process, especially when datasets surpassed the available RAM of the workstation, a two-step approach was adopted. First, substacks of the 3D datasets were generated using a FIJI script, with each substack containing approximately 15% of the central data from the full stack. Subsequently, each substack was stitched to its adjacent counterpart, ensuring optimal overlap based on cross-correlation measures. Using the localization data from each substack post-stitching, the complete 3D stacks were then assembled.

The 3D data visualization was achieved using the Surpass view in Imaris (Version 10.0.1, Bitplane Inc., Zurich, Switzerland). All data processing tasks were executed on a HIVE workstation (ACQUIFER Imaging GmbH, Germany) equipped with dual Intel Xeon Gold 6252 CPUs (2.1 GHz, 24 cores), 1 TB of memory, and an Nvidia RTX A4000 GPU (16 GB GDDR6), operating on Windows Server 2019.

### Image analysis

Images obtained from the Yokogawa microscope were processed and analyzed in MATLAB (2021a, Mathworks, RRID:SCR_001622) using a previously described image analysis pipeline (Bolognin et al., 2019; Monzel et al., 2020).

### Electrochemical detection of catecholamines in neuronal tissues

Commercially available Nafion-coated carbon fiber microelectrodes (World Precision Instruments, CF10-50) were employed to measure the presence of catecholaminergic neurotransmitters (e.g., dopamine) within the assembloids. Amperometric measurements were performed according to the manufacturer’s instructions using a potentiostat (Bio-Logic, VMP3) equipped with a low current module (Bio-Logic) and a custom-made platform (Supplementary Figure 11.D). Electrodes were activated by applying a potential of 1.2V and simultaneous exposure to a 150 mM NaCl solution with a pH of 9.5. All measurements were conducted at a potential of 0.65V. Prior to the measurements, assembloids were washed three times using PBS. Assembloids were subsequently placed onto the custom-made platform, covered in 100 µl of PBS before electrodes were carefully inserted into the tissue using a laboratory jack. Measurements on StrOs alone and StrOs within D30 assembloids were conducted by introducing the electrode 1-2 times into the centre of the neuronal tissue. For the StrOs cultured in pre-used MOs media, the media was in contact with the MOs for 24 hours. Then it was transferred into the wells containing StrOs. StrOs were cultured in the pre-used MO media for 48 hours prior to the measurements. Due to the occasional occurrence of necrotic cores within assembloids electrochemical measurements on D60 assembloids were performed by introducing the electrodes at the border of the StrO. Measurements were conducted at five different locations within the StrO of the assembloid. To exclude any cross-contaminations, electrodes were carefully washed, and background signals were recorded in between the measurement of each assembloid. Measurement results were averaged and unless stated otherwise data was background subtracted.

### Quantitative PCR

The RNeasy Mini Kit (Qiagen, 74106) was used for the total RNA extraction form striatum organoids. The RNA concentration was measured using the Nanodrop 2000c Spectrophotometer (Thermo Fisher Scientific, RRID:SCR_020309). The High-Capacity RNA-to-cDNA^TM^ Kit (Thermo Fisher Scientific, 4387406) was used for the cDNA synthesis. For the quantitative PCR reaction, the Maxima SYBR Green qPCR Master Mix (Thermo Fisher Scientific, K0221) was used with the primers listed in Supplementary Table 4. The Aria Mx Real-Time PCR system (Agilent) was used and the data were extracted from the AriaMx PC software.

### RNA sequencing

RNA was extracted from the assembloids using the RNeasy Mini Kit. Four assembloids were used per condition and from three batches. The RNA samples were shipped with dry ice to Novogene in UK for the RNA sequencing experiment and bioinformatic analysis. The following methodology was used:

### Quality control of reads

In the process of obtaining clean reads, reads containing adapters, higher than 10% undetermined bases and low quality (Qscore of over 50% bases of the read is <=5) were removed.

### Library Construction, Quality Control and Sequencing

Poly-T oligo-attached magnetic beads were used to purify the messenger RNA from the total RNA. After fragmentation, the first strand cDNA was synthesized using random hexamer primers, followed by the second strand cDNA synthesis using dUTP for directional library. To quality control the library, Qubit and real time PCR were used for quantification, while for the size distribution detection bioanalyzer was used. The quantified libraries were combined and sequenced on Illumina platforms, taking into consideration the optimal library concentration and desired data volume.

### Clustering and sequencing

The clustering of the index coded samples was performed according to the manufacturer’s instructions. Following the generation of clusters, the library preparations underwent sequencing using an Illumina platform, resulting in the generation of paired-end reads.

### RNA sequencing Data Analysis

#### Quality control

The initial processing of the raw data (raw reads) in fastq format involved utilizing the fastp software (RRID:SCR_016962). This step involved removing reads that contained adapters, reads with poly-N sequences, and low-quality reads from the raw data, resulting in obtaining clean data (clean reads). Additionally, metrics such as Q20, Q30, and GC content were calculated for the clean data. All subsequent analyses were performed using the high-quality clean data.

#### Reads mapping to the reference genome

The reference genome (hg38) and gene model annotation files were directly downloaded from the genome website. To enable alignment of the paired-end clean reads, an index of the reference genome was constructed using Hisat2 v2.0.5 (RRID:SCR_015530). Subsequently, Hisat2 v2.0.5 was employed as the mapping tool of choice. We specifically chose Hisat2 due to its ability to generate a splice junction database utilizing the gene model annotation file, resulting in improved mapping accuracy compared to non-splice mapping tools.

#### Quantification of gene expression level

To determine the number of reads mapped to each gene, featureCounts v1.5.0-p3 (RRID:SCR_012919) was employed. Subsequently, the Fragments Per Kilobase of transcript sequence per Millions (FPKM) base pairs sequenced for each gene was calculated based on the gene’s length and the count of reads mapped to it. FPKM is a widely utilized method for estimating gene expression levels as it accounts for both the sequencing depth and gene length when considering the reads count. It provides a comprehensive measure of gene expression and is currently the most commonly used approach in this regard.

#### Differential expression analysis

Differential expression analysis was conducted on two conditions/groups using the DESeq2 R package (version 1.20.0, RRID: SCR_015687). DESeq2 utilizes a statistical model based on the negative binomial distribution to determine differential expression in digital gene expression data. The resulting P-values were adjusted using the Benjamini and Hochberg’s method to control the false discovery rate. Genes with an adjusted P-value of less than or equal to 0.05, as determined by DESeq2, were identified as differentially expressed.

Before performing the differential gene expression analysis, the read counts for each sequenced library were adjusted using the edgeR package (version 3.22.5, RRID:SCR_012802) by applying a scaling normalization factor. The differential expression analysis of the two conditions was carried out using the edgeR. The P-values were adjusted using the Benjamini and Hochberg’s method. A corrected P-value threshold of 0.05 and an absolute fold change of 2 were set to determine significantly differential expression.

#### Enrichment analysis of differentially expressed genes

The clusterProfiler (RRID:SCR_016884) R package was used for Gene Ontology (GO) (RRID:SCR_002811) enrichment analysis on the differentially expressed genes, with gene length bias being corrected. GO terms with a corrected P-value < 0.05 were deemed significantly enriched by the differentially expressed genes.

For the analysis of KEGG (RRID:SCR_012773) pathways, which provide insights into the functions and utilities of biological systems, especially based on large-scale molecular datasets, the clusterProfiler R package was employed. The statistical enrichment of differentially expressed genes in KEGG pathways was assessed.

### Single nuclei RNA sequencing

#### Samples processing

Samples were processed and sequenced by Singleron. 10 MOs, 10 StrOs and 4 assembloids per batch form 2 batches, generated from the same wild type (WT) line (201, Supplementary Table 1) were snap frozen in 1.5 ml Eppendorf tubes, and sent to Singleron in dry ice. The culture time point was D30 for assembloids, D50 for MOs and D65 for StrOs. As mentioned before, assembloids were generated by the merging of D20 MOs and D35 StrOs. MOs and StrOs that were cultured separately after D20 and D35, were cultured in the same media of the assembloid condition (optimized co-culture medium). Samples from the two batches were pulled together for nuclei extraction and sequencing by Singleron.

#### Data analysis

Reads were mapped to Homo_sapiens_ensembl_92 genome, and 10X matrices were generated for each sample. To perform the downstream analysis, we used R studio (23.06.1+504 version, RRID:SCR_000432) and R 4.2.2 version (RRID:SCR_001905). Using the Seurat package (version 4.3.0, RRID:SCR_016341), Seurat objects were created for each sample and quality control filtering was performed. For all datasets, cells with >5% mitochondrial genes were filtered out. Additional filtering for cell debris and doublets was performed for each dataset. In MOs data cells with <100 and >1000 genes, in StrOs cells with <100 and >1800 genes and in assembloids cells with <100 and >4000 genes were filtered out. Similar to another study, after filtering, ribosomal and mitochondrial genes were removed from all datasets, as they are considered contamination in the single nuclei RNA sequencing experiments (Khan et al., 2021). After filtering, using the standard Seurat workflow, we analyzed each dataset separately, for identifying clusters specific to each model. LogNormalisation was performed in each dataset, followed by the identification of the 2000 most variable genes (FindVariableFeatures function). Next, data were scaled with the ScaleData function and linear dimensionality reduction was performed with the RunPCA function. 10 PCs were used for the MO dataset clustering and 15 PCs for the StrO and assembloid datasets. Clusters were identified using the FindNeighbors and FindClusters functions, at 0.5 resolution in all datasets. Determination of cell type identity in each cluster was performed with the evaluation of specific cellular markers expression using the DotPlot visualization method.

For identifying differentially expressed genes (DEGs) between assembloids and MOs or StrOs, integration analysis was performed (Butler et al., 2018). After integrating the data, DEG lists using the FindMarkers function were computed, defining the assembloid dataset as ident.1 and the MO or StrO dataset as ident.2. The DEG lists from both comparisons were imported in the Metacore (Clarivate, 2023) online software, were enrichment analysis with FDR threshold > 0.25 and adjusted P.value < 0.05 was performed. Enriched pathways from the “Process Networks” category were extracted. Genes related to the enrichment of the “Development_Neurogenesis_Axonal guidance” pathway were exported, and their expression pattern (up or down regulation) was evaluated. DEGs were also used to assess the expression pattern of genes related to neuronal maturity.

#### Multi-electrode Array

Non-embedded assembloids were used for the electrophysiological analysis using the Axion Micro-electrode array (MEA) system. 48-well MEA plates (Axion, M768-tMEA-48B-5) were first coated with 0.1 mg/ml poly-D-lysine (Sigma-Aldrich, P7886) and incubated in 37°C, 5% CO2 overnight, followed by an 1 hour incubation with 1 mg/ml laminin (Sigma-Aldrich, L2020). Laminin coating was removed, and the plates were washed twice with sterile PBS (Thermo Fisher Scientific, 14190250). Each assembloid was placed in the center of the well on the electrodes and after the media was carefully aspirated, it was left for 2-3 min to dry. Then, 15 μl of Geltrex was added on top of each assembloid and was left in the incubator for 5 min to polymerise. 500 μl of fresh culture medium was added in each well and the plate was kept in the incubator (37°C, 5% CO2) under static conditions. Electrophysiological data were acquired with the Axion Maestro Multiwell 768-channel MEA System (Axion Biosystems) and the Axis software (Axon Biosystems, RRID:SCR_016308). For the analysis of the data the spike lists were exported from the Axis software and the data were processed with MATALB (2021a, Mathworks, RRID:SCR_001622). Plots of the MATLAB exported data were generated using R 4.2.2 vesrion.

#### Data analysis and statistics

Data were analysed usings GraphPad Prism 9.0.0 or R studio (23.06.1+504 version) with R 4.2.2 version. Normality test was performed using the Shapiro test. If not stated otherwise, outlier removal was performed using the ROUT method Q 1% in GraphPad or the Inter-Quartile Range (IQR) proximity rule in R and data were batch normalized to the mean value of each batch. For normally distributed data, two-sided Wilcoxon test or Kruskal-Wallis with Dunn’s multiple comparison test and Benjamini-Hochberg correction was implemented. For normally distributed data, Welch’s t-test or one-way ANOVA with Tukey’s multiple comparison test was performed. Significant P value is represented with asterisks in the order P<0.05 *, P <0.01 **, P <0.001 ***, P <0.0001 ****. Error bars represent mean ± SD.

## Data availability

Raw and processed data that support the findings in this study, as well as scripts used for the analysis of the data are publicly available at this link: https://doi.org/10.17881/4va5-e156 Bulk RNA and single nuclei RNA sequencing data are available on Gene Expression Omnibus (GEO) under the accession codes GSE236458 and GSE241632 respectively.

## Results

### Development of the Midbrain-Striatum assembloid model

The nigrostriatal pathway is characterised by dopaminergic neuronal projections from the SNpc to the putamen and caudate of the dorsal striatum. To recapitulate this pathway *in vitro*, we generated a 3D *in vitro* model of midbrain and striatum organoids. MOs were generated based on our previously published protocols (Monzel et al., 2017; Nickels et al., 2020). For the generation of StrO different culture conditions were tested (RA, SR and C4) and compared to the recently published protocol (named C3) (Miura et al., 2020) with minor changes (Supplementary Figure 2). The purpose of this optimisation on the StrOs protocol generation was to reduce the lengthy culture time of 50-80 days to 35-50 days while preserving the mature population of medium spiny neurons (MSNs). For the conditions RA, SR and C4 organoids were cultured for 35 and 50 days, while for the C3 organoids were cultured until D50 (the earliest time point described in (Miura et al., 2020)). In C4, the differentiation conditions are similar to C3 (Miura et al., 2020), but in the maturation phase (D17 to D35), DHA was excluded as it is susceptible to oxidation and to avoid the use of ethanol (the solvent for DHA) in the medium. The same medium was used until D50 without the addition of DAPT at D42 (Supplementary Figure 2.A). DHA was also excluded from the C3 (Supplementary Figure 2.B). For conditions RA and SR (Supplementary Figure 2.C, D) the previously published protocol which described the generation of ventral forebrain, subpallium-like organoids (Sloan et al., 2018), was adapted to shift the differentiation towards the LGE region and eventually to the dorsal striatum. In both conditions, SAG was added as it is neuroprotective and very important for neurodevelopment (Nguyen et al., 2021), while activin A has been shown to assist the differentiation of striatal projection neurons (Arber et al., 2015). As was demonstrated by Miura and colleagues (Miura et al., 2020), SR11237 was used to stimulate the RXRG receptor (SR condition), which has high expression in the development of the striatum as shown in the human brain transcriptome (HBT) database. Similarly, we also wanted to test the effects of RA supplementation (RA condition) as data in the HBT database show that RARB is also highly expressed in the first days of striatum development.

Gene expression analysis with qPCR for genes specific for the telencephalon and the LGE were used to confirm the differentiation of the organoids towards the dorsal striatum (Supplementary Figure 3.A). Although the forebrain marker Forkhead box protein G1 (*FOXG1*) was evenly expressed in all conditions, there were some differences in the expression of LGE progenitor genes. Achaete-Scute Family BHLH Transcription Factor 1 (*ASCL1*) showed a tendency of higher expression in the RA and SR conditions in both time points, indicating a slower differentiation of cells with higher number of progenitors at D50 of culture. In C4 at D35, *ASCL1* had similar expression with RA and SR conditions, but at D50 the expression was lower and similar to C3. Genetic-Screened Homeobox 2 (*GSX2*) is an essential transcription factor in LGE progenitors, that assists their differentiation towards neurons and glia cells in the dorsal striatum (Roychoudhury et al., 2020). In the here investigated conditions we show that *GSX2* is mainly expressed in the C4 and C3 at D50. The Forkhead Box P1 (*FOXP1*) and Forkhead Box P2 (*FOXP2*) genes are both transcription factors that are highly expressed in the striatum and are important for the development of MSNs (Fong et al., 2018). The expression of *FOXP1* was similar in all conditions, but *FOXP2* had higher expression in C4 and C3 at D50, indicating a better differentiation of the neurons towards MSN fate. In addition COUP-TF-interacting protein 2 (*CTIP2*), another crucial transcription factor for the differentiation of MSNs (Arlotta et al., 2008), showed higher expression in the organoids of C4 and C3. Expression of *NKX2.1* and *OTX2* genes are important for the development of the medial ganglionic eminence (MGE) (Sandberg et al., 2018). The low expression of these genes in C3 and C4 further validates a LGE and dorsal striatum identity (Supplementary Figure 3.B). The presence of mature MSNs in the organoids was evaluated by the expression of the MSN specific marker Dopamine and cAMP-Regulated Neuronal Phosphoprotein 32 (*DARPP32*) and the dopaminergic receptors 1 and 2 (*DRD1* and *DRD2*). We observed that C4 had noticeable higher expression of *DARPP32*, while the expression of *DRD1* and *DRD2* was similar between C3 and C4 (Supplementary Figure 3.C). Similar patterns were observed in the protein levels, where DARPP32 abundance was similar at D50 for conditions C3 and C4, DRD1 was higher in C3 and C4 at D50 and there were no differences in the levels of DRD2 between all conditions. Glutamic acid decarboxylase 65-kilodalton isoform (GAD65), important enzyme for the synthesis of GABA, was higher in C3. Overall, these results indicate that C4 and C3 have similar differentiation capacity towards dorsal striatum identity (Supplementary Figure 3.D-G). C4 seems to result in progenitor rich striatum organoids at D35 with differentiated characteristics at D50.

Since MOs and StrOs were differentiated independently for their identity specification, the generation of the assembloid model firstly required the optimisation of a suitable co-culture medium. Assembloids were generated between D20 of MOs derived from a GFP-expressing cell line and D35 or D50 of StrOs (Supplementary Figure 4.A), from the four StrO conditions described before (Supplementary Figure 2). Four different medium conditions were tested to ensure the optimal development and identity specificity of both organoids (Supplementary Figure 4.B). First, we evaluated the development of the dopaminergic neurons in GFP-expressing MOs cultured for up to 20 days by staining for TH and MAP2. MOs cultured in NM and NMpl media showed significant depletion of dopaminergic neurons, while the organoids in NM++ and N2B27++ medium did not show significant differences from the control condition (MOs standard media, see Midbrain organoids section in Methods) (Supplementary Figure 4.C). Then, the four different media were tested in assembloids cultured for 20 and 35 days to assess the impact on both dopaminergic neurons (TH staining) and MSNs (DARPP32 staining). Similar to the observations in MOs, assembloids cultured with NM++ and N2B27++ media, showed high levels of TH neurons in both time points of assembloid culture, specifically in C4 and C3 (Supplementary Figure 5). Since these two conditions were more suitable for the development of TH-positive neurons and the N2B27++ was slightly more favourable for the DARPP32 positive neurons in C4 (Supplementary Figure 6), we used N2B27++ as the optimal co-culture medium for the assembloids (Figure 1.A). Overall, based on these data, in order to reduce the time of cultures and achieve the desired differentiation capacity in assembloids, StrOs generated with the strategy C4 at D35 of differentiation were used for the generation of assembloids in the subsequent experiments.

**Figure 1:**
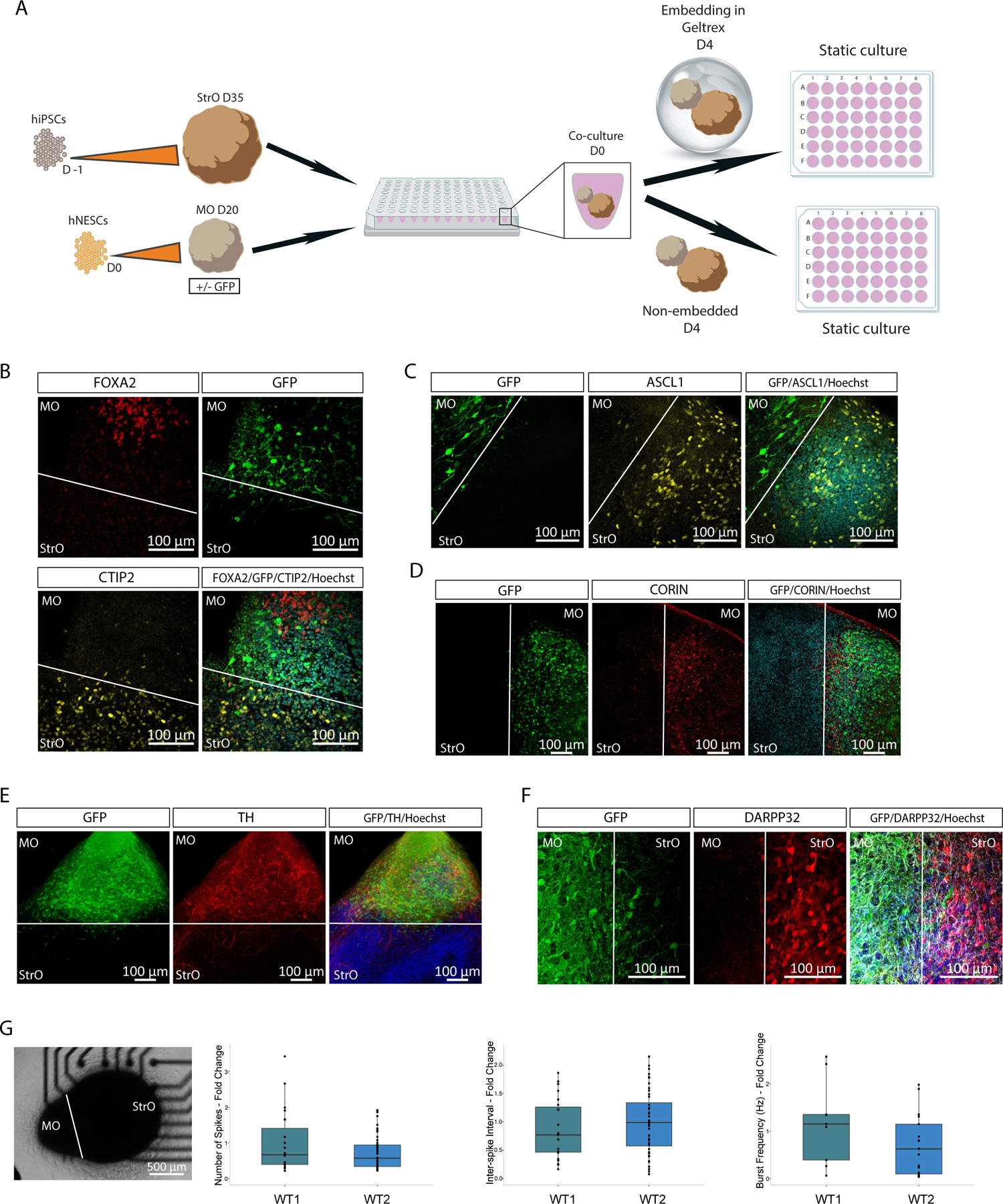
Generation of midbrain-striatum assembloid model with identity specificity. **A.** Schematic representation of the assembloid model generation. **B.** Representative confocal image of a 70 μm assembloid section immunostained with Hoechst, FOXA2 visible in the MO side and CTIP2 visible in the StrO side of the assembloid. GFP fluorescence is intrinsic in the midbrain part of the assembloids. **C.** Representative confocal image of a 70 μm assembloid section immunostained with Hoechst and ASCL1 visible in the StrO side of the assembloid. GFP fluorescence is intrinsic in the MO part of the assembloids. **D.** Representative confocal image of a 70 μm assembloid section immunostained with Hoechst and CORIN visible in the MO side of the assembloid. GFP fluorescence is intrinsic in the midbrain part of the assembloids. **E.** Representative confocal image of a 70 μm assembloid section immunostained with Hoechst and TH visible in the MO side of the assembloid. GFP fluorescence is intrinsic in the midbrain part of the assembloids. **F.** Representative confocal image of a 70 μm assembloid section immunostained with Hoechst and DARPP32 visible in the StrO side of the assembloid. GFP fluorescence is intrinsic in the midbrain part of the assembloids. **G.** Representative brightfield image of an assembloid in a well of the 48-well MEA plate. Plots showing the quantification of the Number of Spikes, Inter-spike Interval in seconds and the Burst Frequency in Hz between assembloids generated from two independent human WT cell lines. The data in the plots represent recordings of individual assembloids for the culture periods D32 to D43, from 3-4 batches. Batch correction was applied by normalising each value to the mean of the values for each batch. Outliers were calculated in GraphPad Prism using the ROUT method Q 1%. Two-sided Wilcoxon test was performed in R 4.2.2.

### Midbrain and striatum specific identity in the assembloid model

Characterisation was performed on assembloids cultured for 30 days. Microscopy assessment with immunofluorescence staining showed that the midbrain and the striatum organoids retain their identity in the assembloid model (Figure 1.B-F). Midbrain progenitors positive for FOXA2 and CORIN were identified only in the midbrain side of the assembloid, while striatal progenitors positive for ASCL1 and CTIP2 appear only in the striatum side (Figure 1.B, C, D). Additionally, mature neurons of midbrain and striatal identity were observed with the positive immunofluorescence staining of TH and DARPP32, respectively (Figure 1.E, F). To validate that the assembloid model exhibits neuronal activity, electrophysiological measurement with MEA were performed on assembloids generated from two independent WT cell lines. Analysis of the MEA recordings showed that assembloids from both lines display similar electrophysiological activity with no differences in the number of spikes, the interspike interval and the burst frequency (Figure 1.G).

To confirm the qualitative observations, we performed single nuclei RNA sequencing on pooled assembloids from two batches, generated from the WT 1 cell line (see Supplementary Table 1) and cultured for 30 days. From the same batches, we also sequenced MOs and StrOs that after D20 and D35 respectively (time points used for assembloid generation) were cultured independently for 30 more days in the same assembloid co-culture medium. Datasets from the three models were analysed separately following the standard Seurat workflow. For the assembloid model, eight different clusters were identified and visualised with UMAP (Figure 2.A). The identity of each cluster was defined based on the expression of cellular specific markers from gene lists identified by literature (Bhaduri et al., 2020; Kamath et al., 2022; La Manno et al., 2016) and the PanglaoDB database (Supplementary Figure 7). Calculation of the percentage of each cellular identity in the assembloid model revealed the presence of 5% A10 dopaminergic neurons (DANs(A10)), 6% of GABAergic neurons, 7% progenitors of the LGE (LGE Prog), 14% of radial glia cells (RGCs), 15% of medium spiny GABAergic neurons (MSNs), 16% of non-defined progenitors (Progenitors), 17% of A9 dopaminergic neurons (DANs(A9)) and 21% young neurons (yNeurons) (Figure 2.B). Hierarchical clustering of the cellular populations based on their top 500 variable genes, demonstrated the similar transcriptomic identity of the dopaminergic neuronal clusters, followed by the other mature neuronal populations of MSNs and GABAergic neurons, while young neurons and progenitor cells cluster together (Figure 2.C). The identity similarity between clusters is further demonstrated by the Spearman’s correlation matrix (Figure 2.D). This illustrates the close correlation of the progenitor clusters (LGE Prog, RGCs and Progenitors) that weakly correlate with the more mature populations. Additionally, there is a weak correlation between the A9 and A10 DANs, indicating their distinct genetic signature. Although A10 DANs exhibit low correlation with all the clusters, their slightly better correlation with the RGCs and Progenitors clusters, suggests a more immature identity. On the other hand, A9 DANs correlate with the more mature clusters (GABAergic Neur and MSNs), while showing a weak correlation with the progenitor clusters. Finally, yNeuron cluster exhibits an unidentified young neuronal identity, correlating with Progenitors, DANs(A9) and GABAergic neurons, but with highest correlation to the MSN cluster.

**Figure 2:**
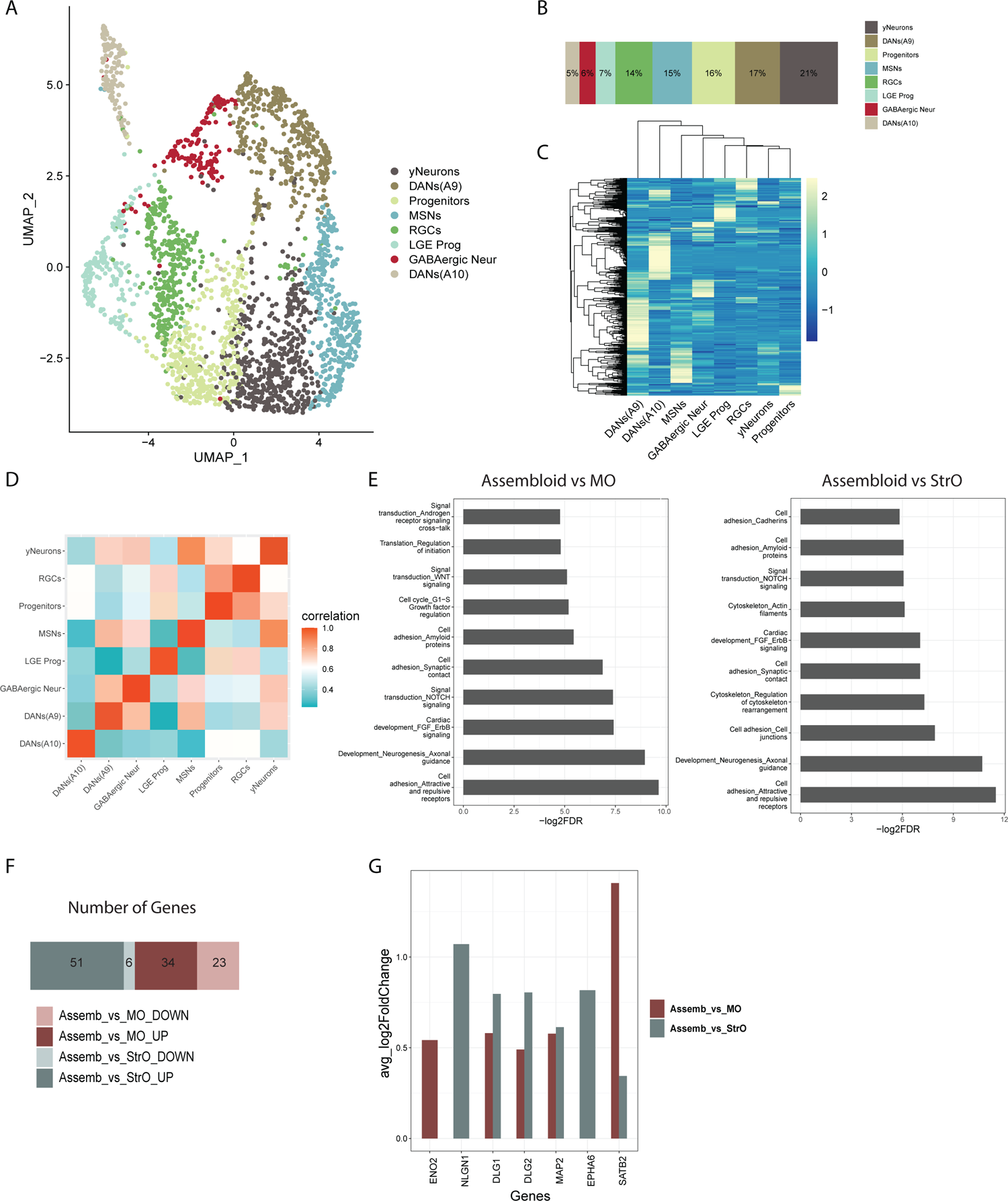
Single nuclei RNA sequencing analysis in the assembloid model. **A.** UMAP embedding of the different cellular clusters in assembloids. **B.** Percentage of the cellular composition in the assembloid model. **C**. Unsupervised hierarchical clustering of cell clusters, using the average expression with Z-score normalisation of the top 500 most variable genes. **D.** Spearman’s correlation between the different cell types in assembloids. **E.** Enriched pathways identified by Metacore using the DEGs between assembloid and MO, and between assembloid and StrO, after integration of the three datasets with the Seurat workflow. **F.** Plot showing the number of genes that were identified in the enrichment of the “Developmental Neurogenesis and Axonal guidance” pathway in both comparisons Assembloid vs MO and Assembloid vs StrO. **G.** Barplot showing the upregulation of neuronal maturity related genes in both DEG lists, of Assembloid vs MO and Assembloid vs StrO.

The same analysis was performed for the MO and StrO datasets, with the presence of cell clusters more specific to the midbrain and striatum identity respectively. MOs are comprised of dopaminergic neuronal clusters (DANs and DANs2, 28% and 3% respectively) but also GABAergic neurons (33%), general neurons with no specific identity (Neurons, 14%), radial glia cells (RGCs, 20%) and a small proportion of oligodendrocyte precursor cells (OPCs, 1%) (Supplementary Figure 8.A, B). Heatmap of the 500 most variable genes, shows the clustering of RGCs in the middle, giving rise to the DANs, GABAergic neurons and Neurons clusters, while DANs2 and OPCs cluster separately, showing a more specified identity (Supplementary Figure 8.C). In Spearman’s correlation, DANs2 cluster shows a more mature identity with a very weak correlation to the progenitor RGCs cluster and a stronger correlation with the GABAergic Neur and Neurons clusters. On the other hand, DANs cluster shows a less mature identity (Supplementary Figure 8.D). The clustering was confirmed by the expression of different cellular markers (Supplementary Figure 8.E-J).

StrOs show a striatum specific cellular composition with 42% MSNs, 39% GABAergic interneurons, 10% LGE progenitors, 8% Neural progenitors and 1% of more general Telencephalic progenitors (Supplementary Figure 9.A-B). Similarly here, hierarchical clustering of the 500 most variable genes shows the close association of the progenitor clusters, followed but the more mature populations of MSNs and GABAergic interneurons (Supplementary Figure 9.C). The maturity of the MSNs is further confirmed by the Spearman’s correlation matrix, with their weak correlation to LGE and Telencephalic progenitors, and their strong correlation to the GABAergic interneurons (Supplementary Figure 9.D). The identity of each cellular cluster was also here validated by the expression of cellular specific markers (Supplementary Figure 9.E-K).

To be able to compare the three datasets and evaluate the genes that are differentially expressed between assembloids and the individual organoids, we performed integration analysis using the Seurat workflow (Butler et al., 2018). A clear shift of cellular populations from the MOs and StrOs into the assembloid model is visible in the UMAP plot of the integrated object based on the clusters identified separately in each model (Supplementary Figure 10.A). The progenitor clusters in all models appear in the upper part of the UMAP. DANs(A10) population clusters closer to the RGCs, illustrating once more their immature identity. Between the progenitor and mature clusters appears an intermediate more general cluster of progenitors in the assembloids model (Progenitors). This progenitor cluster seems to evolve into neuronal identity cells, given its close association with the yNeurons cluster. DANs cluster, that has a more immature identity in MOs, was not present in the assembloid model, while the general neuronal cluster in MOs, seems to have been shifted towards the DANs(A9) identity in assembloids. The MSNs cluster from StrOs, was preserved in assembloids. The strong correlation between GABAergic interneurons and MSNs is likely the reason why this cluster is not observable in the assembloid model (Supplementary Figure 10.B). Additionally, a small number of OPCs is only present in MOs. Given their correlation with neuronal cluster, this cluster probably consists of cells with mixed identity (Supplementary Figure 10.B).

Enrichment analysis using the DEGs of assembloids-MOs and assembloids-StrOs, revealed the enrichment of several processes related to synaptic contact, cell adhesion but also neurogenesis and axonal guidance (Figure 2.E). Concerning the developmental neurogenesis and axonal guidance pathway, which was the second most enriched pathway in both comparisons, we observed that the majority of responsible genes for enriching this pathway are upregulated in the assembloid model compared to the MO and StrO models (Figure 2.F). Additionally, we evaluated the expression of genes related to neuronal maturity and identified six genes, all showing upregulation in the assembloid model (Figure 2.G). Finally, genes related to cellular and oxidative stress were found downregulated in assembloids compared to MOs and StrOs (Supplementary Figure 10.C, Supplementary Table 5).

### Midbrain-striatum assembloid model resembles the nigrostriatal pathway connectivity

For validating the existence of nigrostriatal pathway connectivity, we examined the presence of dopaminergic projections from the midbrain into the striatum in the assembloid model. First, to confirm that TH+ signal in the striatum of the assembloids originates from the MOs TH+ neurons innervation, we examined the presence of TH+ neurons in GFP-expressing MOs and StrOs cultured separately in the assembloid co-culture media for the same culture period as the assembloids (D30 assembloids, D50 MOs and D65 StrOs). We observed that MOs develop a high number of TH+ neurons, while the StrOs have almost no TH+ signal (Supplementary Figure 11.A). Next, for visualising the TH+ neurite projections from the midbrain to the striatum in the assembloid model, we performed whole mount imaging of D30 assembloids containing GFP-expressing MOs. We were able to identify neurons that have TH+/GFP+ soma, indicating their midbrain identity, with TH+ axons projecting towards the striatum side of the assembloid (Figure 3.A, Supplementary Figure 11.B, C).

**Figure 3:**
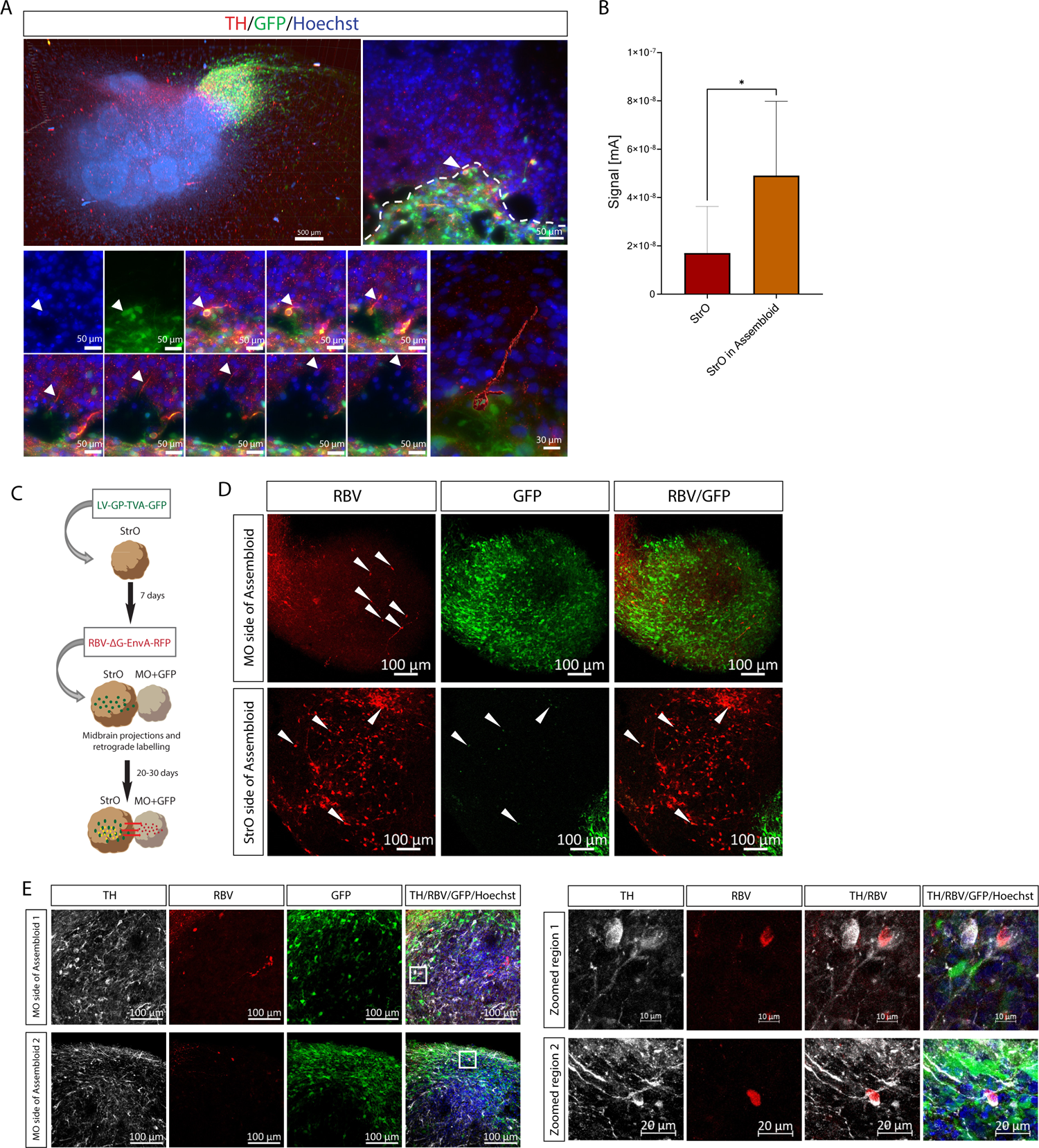
Midbrain-Striatum assembloids develop nigrostriatal pathway connectivity. **A.** Representative microscopic images of a whole assembloid (4X objective) and of ROI, observed with fluorescence microscopy (25X objective). The assembloid was immunostained with Hoechst and TH. The ROI (right panel) shows the GFP+/TH+ neuron’s soma in the MO-GFP side of the assembloid with TH+ projection towards the striatum side in the different planes along the Z stack (Z.190-280). White arrowheads show the progression of the TH+ projection in images from the different planes. 3D reconstruction of the neuron across the planes shows the complete TH+ neuronal projection. **B.** Bar pot showing the electrochemical measurements in tissue in StrOs and in the StrO side of the assembloid model at D30. Welch’s t-test was performed, StOs n=8, StrOs in assembloid n=9, where n is the average measurements in one organoid or assembloid, for three batches, generated from the same line (390, see Supplementary Table 1). Error bars represent mean ± SD. Data were plotted in GraphPad Prism 9.0.0. *p<0.05, **p<0.01, ***p<0.001. **C.** Schematic representation of the Rabies monosynaptic tracing experiments in midbrain-striatum assembloids. **D.** Representative confocal image of 70 μm assembloid section showing the GFP and RFP positive cells from the LV-GP-TVA-GFP and RBV-ΔG-EnvA-RFP infections respectively. The GFP signal in the StrO side of the assembloid is coming from the LV-GP-TVA-GFP infection, while GFP fluorescence in the MO side of the assembloid is cell intrinsic. White arrowheads indicate the RFP positive signal in the MO side, and the RFP/GFP positive signal in the StrO side of the assembloid. **E.** Representative confocal images of 70 μm assembloids sections showing the MO sides of the assembloids with GFP and RFP positive cells from the cell intrinsic GFP expression and RBV-ΔG-EnvA-RFP infections respectively and immunostained with Hoechst and TH. Zoomed in regions (indicated by the white squares) show the TH+/RFP+ colocalization in the midbrain side of the assembloids.

Nigrostriatal pathway connectivity and functionality in the assembloid model were further investigated. Catecholamine levels were assessed by inserting a Nafion-coated carbon electrode into the neuronal tissues (Supplementary Figure 11.D). We have previously shown that electrochemical monitoring of catecholamine levels in the supernatant of MOs can provide valuable insights into the presence of dopamine (DA), due to neglectable levels of the interfering cationic catecholamines norepinephrine and epinephrine within the system (Zanetti et al., 2021). As expected, we observed higher levels of catecholamines in the MO side of the assembloid compared to the StrO side (Supplementary Figure 11.E). No differences in StrO and StrO cultured in MO-conditioned media were detected, indicating no significant catecholamine or DA uptake from the medium (Supplementary Figure 11.F). Finally, we measured catecholamine levels in StrOs alone and StrOs in the assembloid model. The results showed that the StrOs in the assembloid model displayed higher signals compared to StrOs cultured alone (Figure 3.B), suggesting the active secretion of the catecholamine DA from midbrain dopaminergic neurons that are projecting to the striatum in the assembloid model.

To further determine whether neurons of MO and StrO are connected through active synapses in the assembloid model, we established a rabies virus-based retrograde monosynaptic tracing system. This system allows tracing of monosynaptic connections between neurons, using a lentiviral vector (Miyamichi et al., 2011) that expresses histone 2B-tagged green fluorescent protein (GFP), the avian tumor virus A (TVA) receptor (for selective infection by EnvA-pseudotyped rabies virus), and rabies virus envelope spike glycoprotein (GP) (to allow for rabies virus transsynaptic spreading) (LV-GP-TVA-GFP), in combination with a recombinant G-deleted (ΔG) rabies viral vector, pseudotyped with EnvA envelope protein (selectively binding to TVA), and red fluorescent protein (RFP)-tagged (RBV-ΔG-EnvA-RFP). LV-GP-TVA-GFP-transduced neurons (starter neurons) are identified by nuclear GFP expression. Following infection with the RBV-ΔG-EnvA-RFP, the double-transduced starter neurons are traced by co-expression of nuclear GFP and RFP, and because of GP expression, allow ΔG-rabies virus to form infectious particles in their cytoplasm. Neurons laying presynaptic inputs on the targeted starter neurons, are rendered RFP+ due to the selective retrograde transmission of rabies virus across active synapses (target neurons) (Ugolini, 1995). Therefore, target neurons are RFP+/GFP-whereas starter neurons are RFP+/GFP+. Only first-order synapses are traced because GP expression is confined to the starter neurons, not allowing for further propagation of ΔG-rabies virus (Etessami et al., 2011; Grealish et al., 2015; Osakada & Callaway, 2013).

StrOs at D35 of culture were transduced with LV-GP-TVA-GFP expressing the H2B-GFP fusion protein under the human Synapsin promoter. One week later, transduced StrOs were merged with MOs expressing GFP to generate assembloids, and together they were infected with RBV-ΔG-EnvA-RFP. The targeted starter neurons in the StrO were RFP+/GFP+. One week later, the media was changed and the assembloids were further cultured for up to 30 days (Figure 3.C). Sections from fixed assembloids were analysed with confocal microscopy for RFP and GFP expression (Figure 3.D). Starter neurons (RFP+/GFP+, arrowheads in Figure 3.D StrO side) as well as target neurons (RFP+/GFP-, arrowheads in Figure 3.D MO side) were detected in the striatum side of the assembloid, demonstrating high connectivity of the nearby neurons with the starter neurons in the striatum organoid. Notably, RFP+ (arrowheads in Figure 3.D MO side) target neurons were additionally present more distantly in the MO compartment, indicating active synaptic connectivity between the two organoids on the assembloid level. TH-immunostaining revealed TH+/RFP+ target neurons in the MO side of the assembloid, suggesting the presence of synapse connectivity between dopaminergic neurons in the MO and starter neurons in the StrO (Figure 3.E). Target neurons (RFP+/GFP-) presence was also confirmed in assembloids where MOs were derived from iPSCs that initially did not express GFP (Supplementary Figure 11.G).

### Doxycycline inducible Progerin overexpression in the assembloid system

For the induction of aging phenotypes in the assembloid model we used an iPSC line that was genetically engineered with a transgene containing the modified *LMNA* gene for the transcription of Progerin, under the control of the Tet-On system, where doxycycline supplementation is needed for the induction of the transgene expression (Gabassi et al., *In Preparation*). For better identification of the Progerin+ cells, Progerin is co-expressed with the GFP fluorescent protein. This line was used to generate assembloids (Figure 4.A), and optimal concentration of doxycycline supplementation was determined by testing three different concentrations (1, 2 and 4 ng/μl). Assembloids were treated with doxycycline from D4 to D30 of culture. At D30, three assembloids per condition and from three batches were used in flow cytometry to measure the amount of GFP positive cells. Approximately 50% of cells were GFP-positive in assembloids treated with the highest concentration of doxycycline (4 ng/μl), which was significantly higher than in the other conditions (Figure 4.B). Similar to what was observed for the GFP levels, Progerin levels, validated by Western blot using the LMNA antibody (see Supplementary Table 2), were significantly higher in the 4 ng/μl doxycycline concentration (Figure 4.C). To evaluate the viability of assembloids treated with doxycycline we performed ATP and LDH assays (Supplementary Figure 12). Here, we also used assembloids generated from the isogenic line (without the Progerin transgene, referred to as WT assembloids) as a control. In the LDH assay, we show significant higher levels of LDH in the 4 ng/μl doxycycline treatment (WT_4_Dox) which indicates that doxycycline could cause some cytotoxicity in the model. However, we noticed a different pattern of LDH levels in the assembloids with the Progerin transgene (Progerin assembloids), with no significant differences between the untreaded (Progerin_0_Dox) and the 4 ng/μl doxycycline treated assembloids (Progerin_4_Dox) (Supplementary Figure 12.A). In the ATP assay, we detected no differences between the conditions, indicating no changes in the viability and metabolic activity of the model when treated with doxycycline (Supplementary Figure 12.B). These findings suggest that 4 ng/μl doxycycline concentration is acceptable for generating a mosaic Progerin-overexpressing assembloid model.

**Figure 4:**
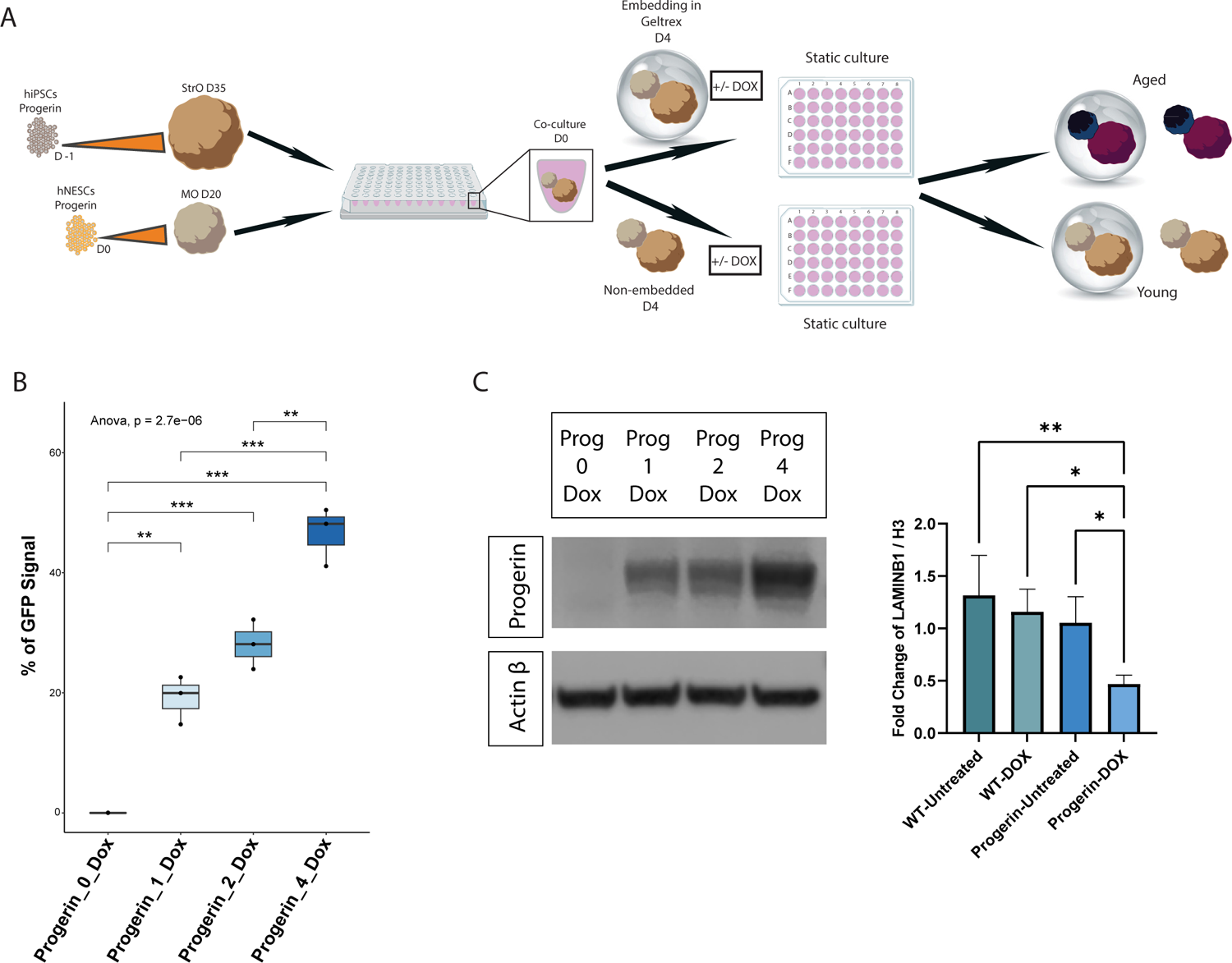
Optimisation of the Progerin-overexpression in the assembloid model. **A.** Schematic representation of the assembloid model generation using the Progerin cell line for the inducible overexpression of Progerin after doxycycline supplementation. **B.** FACs data showing the % of the GFP positive signal measured in live cells of dissociated assembloids, treated with different doxycycline concentrations (Progerin_0_Dox = Untreated, Progerin_1_Dox = 1 ng/μl, Progerin_2_Dox = 2 ng/μl, Progerin_4_Dox = 4 ng/μl). One-way ANOVA, with Tukey’s multiple comparison test was performed in R 4.2.2. For all conditions n = 3 with each point representing the average of two technical replicates per batch for 3 batches. *p<0.05, **p<0.01, ***p<0.001. **C.** Western blot for Progerin protein levels in assembloids treated with the different doxycycline concentrations. One-way ANOVA, with Tukey’s multiple comparison test was performed in R 4.2.2. Error bars represent mean ± SD. For all conditions n =3 with each point representing 3-4 polled assembloids per batch, for 4 batches. *p<0.05, **p<0.01, ***p<0.001. Batch correction was applied by normalising each value to the mean of the values for each batch. Outliers were calculated in GraphPad Prism using the ROUT method Q 1%.

### Progerin-overexpressing assembloids show aging characteristics

To determine whether neurons that express Progerin in the assembloids acquire aging characteristics, we stained assembloid sections from D30 and D60 assembloids treated with 4 ng/μl doxycycline for aging-associated markers (Figure 5). Colocalization of H2AX and 53BP1 positive foci is a marker of DNA double-strand breaks. We were able to see that D60 assembloids had significantly higher levels of H2AX/53BP1 positive foci in the Progerin-expressing cells (marked as GFP+) compared to the non-Progerin-expressing cells (Figure 5.A). Similarly, aging-associated markers such as p21, p16 and p53 were all significantly elevated not only in the Progerin-expressing cells, but more specifically in Progerin-expressing neurons (MAP2/GFP-double positive) in D60 assembloids (Figure 5.B, C, D).

**Figure 5:**
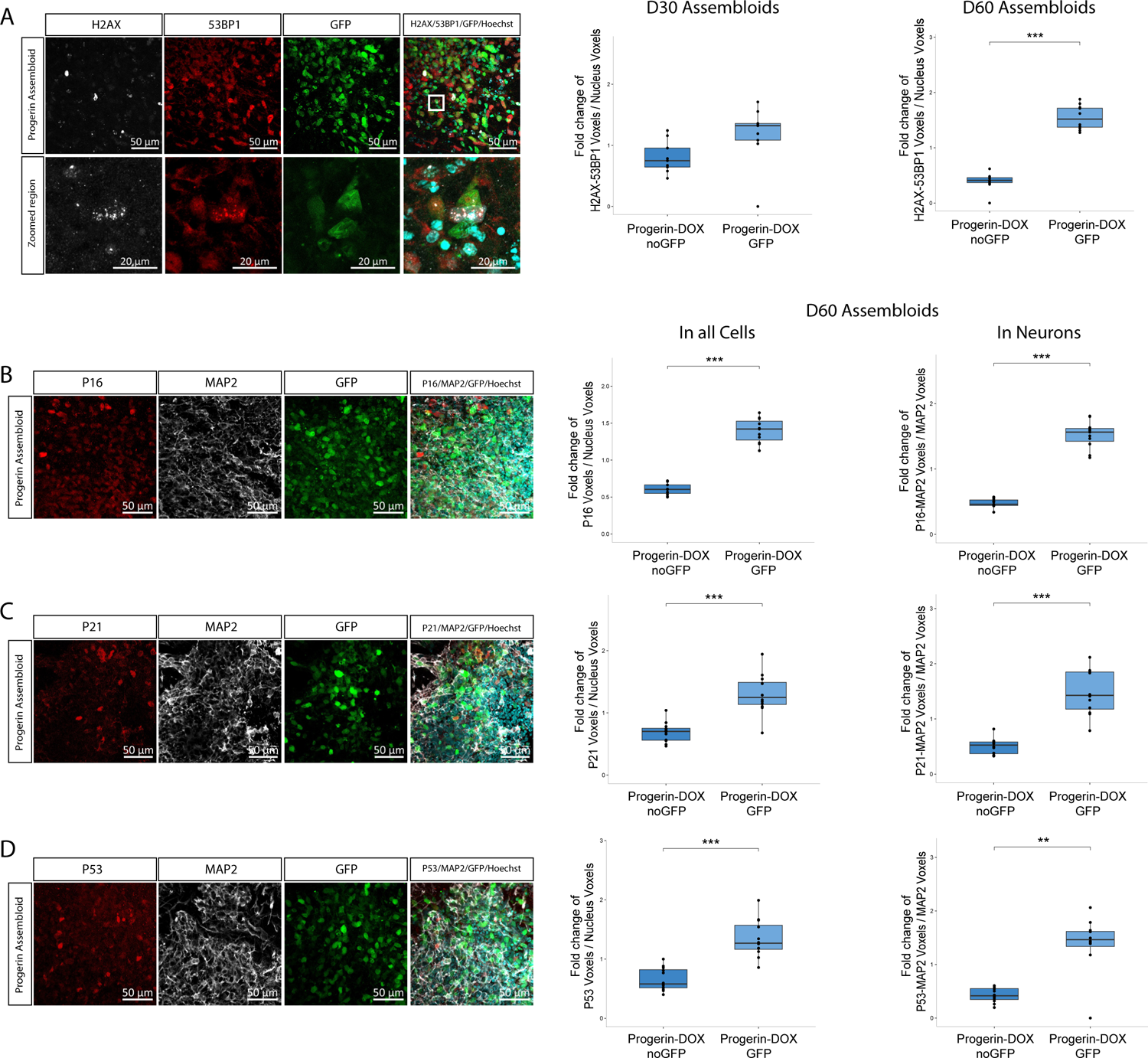
Progerin-overexpressing cells with aging characteristics in the assembloid model. **A.** Immunofluorescence staining quantification of the H2AX and 53BP1 positive nuclear foci voxels normalised to the total nucleus voxels in 70 μm Progerin-overexpressing assembloid sections from D30 and D60 cultures. For D30 data, Welch’s t-test was performed with n = 9 for both conditions where each point represents one section per assembloid per batch for 3 batches. For D60 data two-sided Wilcoxon test was performed with n = 12 for both conditions where each point represents one section per assembloid per batch for 4 batches. *p<0.05, **p<0.01, ***p<0.001. **B.** Immunofluorescence staining quantification of the P16 voxels normalised to the total nucleus voxels, and P16 and MAP2 double positive voxels normalised to the total MAP2 voxels in 70 μm Progerin-overexpressing assembloid sections from D60 cultures. Two-sided Wilcoxon test for both plots was performed, with n = 11 where each point represents one section per assembloid per batch for 4 batches. *p<0.05, **p<0.01, ***p<0.001. **C.** Immunofluorescence staining quantification of the P21 voxels normalised to the total nucleus voxels, and P21 and MAP2 double positive voxels normalised to the total MAP2 voxels in 70 μm Progerin-overexpressing assembloid sections from D60 cultures. Welch’s t-test and two-sided Wilcoxon test was performed respectively, with n = 12 for both conditions where each point represents one section per assembloid per batch for 4 batches. *p<0.05, **p<0.01, ***p<0.001. **D.** Immunofluorescence staining quantification of the P53 voxels normalised to the total nucleus voxels, and P53 and MAP2 double positive voxels normalised to the total MAP2 voxels in 70 μm Progerin-overexpressing assembloid sections from D60 cultures. Welch’s t-test and two-sided Wilcoxon test was performed respectively, with n = 12 for both conditions where each point represents one section per assembloid per batch for 4 batches. *p<0.05, **p<0.01, ***p<0.001. In all plots batch correction was applied by normalising each value to the mean of the values for each batch. Outlier removal was performed based on the Inter-Quartile Range (IQR) proximity rule. Data were plotted in R 4.2.2.

Next, we explored the impact of the Progerin-expressing cells on the whole assembloid model. For that, we performed transcriptomic analysis with bulk RNA sequencing on assembloids generated from the Progerin-expressing cell line and its respective isogenic control. In both cases, we used the doxycycline treated (WT_DOX, PROG_DOX) and the untreated conditions (WT_UNTR, PROG_UNTR). Principle component analysis (PCA) plot shows the clustering of the samples based on their transcriptomic similarity. In this case, all the Progerin-expressing (PROG_DOX) samples were clustered completely separately from the rest of the samples (Figure 6.A). Similarly, a heatmap of the DEGs shows that the Progerin-expressing samples have a distinct expression pattern (Supplementary Figure 13.A). Additionally, the number of DEGs of assembloids with Progerin overexpression (PROG_DOX group) versus the control groups (PROG_UNTR, WT_UNTR, WT_DOX) was 2 or 3-fold higher (Supplementary Figure 13.B). These data demonstrated that assembloids with Progerin overexpression acquire a distinct transcriptomic profile. To explore possible aging-related transcriptomic changes, relevant to the human brain in the Progerin-overexpressing assembloids, we pooled the DEG lists from the comparisons between the Progerin-overexpressing assembloids with the controls (PROG_DOXvsPROG_UNTR, PROG_DOXvsWT_UNTR and PROG_DOXvsWT_DOX) and we compared them to the differentially expressed genes of aged human post-mortem brain transcriptomic data from two studies (Berchtold et al., 2008; González-Velasco et al., 2020). For these comparison, we extracted the significant DEGs (adj. pvalue < 0.05) that are commonly up or downregulated in the post-mortem and assembloid datasets (Extended data 1-4). From these genes we focused on the ones with a log2FoldChange higher than 2. Additionally, we included the expression of *PNOC* (foldchange = −1.34) and *GMPR* (foldchange = 1.72) genes. The expression pattern of these two genes is best associated with aging, according to González-Velasco and colleagues (González-Velasco et al., 2020) (Figure 6.B). The aging associate genes *VIP, SST, NRGN, NETO1, KCNAB1, GRM4, GABRA4, DRD2, DRD1, DLX6, CRH, COBL, CHCHD2, IER3, GMPR, COL21A1* and *CHI3L1* were all significantly differentially expressed.

**Figure 6:**
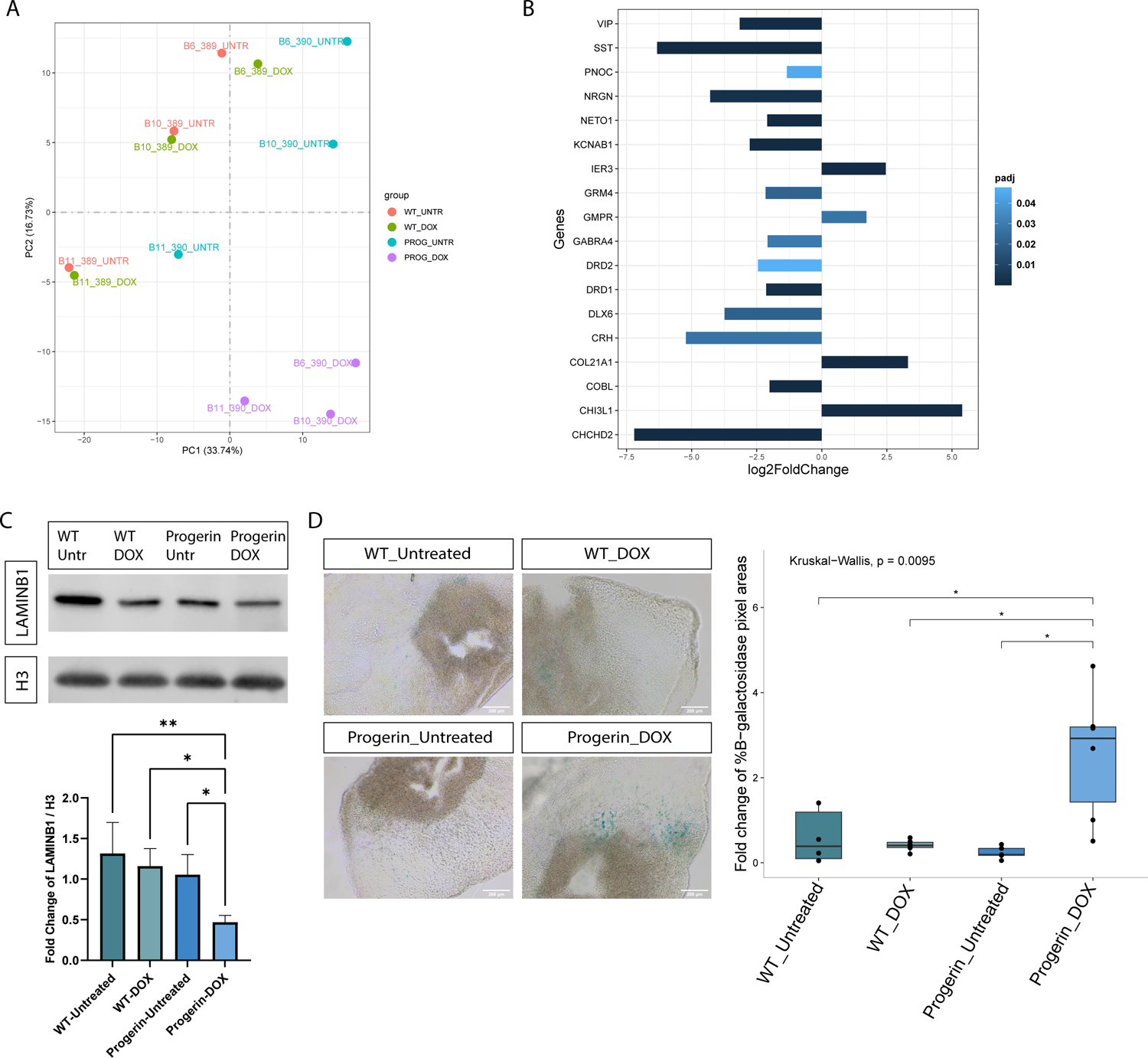
Evident aging phenotype in the Progerin-overexpressing assembloid model. **A.** PCA plot of the two first principal components on the gene expression value (FPKM) of all samples. Each sample represents data from 4 pooled assembloids from one batch. **B.** Plot showing the log2 fold change of significant differentially expressed genes between Progerin-overexpressing assembloids (PROG_DOX) and control (WT_UNTR, WT_DOX, PROG_UNTR) samples. This list of genes was extracted after the comparison of the assembloid data with post mortem human brain data (Berchtold et al., 2008; González-Velasco et al., 2020). **C.** Western blot showing the protein levels of LAMINB1 normalised to H3 housekeeping protein and batch corrected by normalising to the mean of the values for each batch. Outliers were calculated in GraphPad Prism using the ROUT method Q 1%. One-way ANOVA, with Tukey’s multiple comparison test was performed. For all conditions n = 4 with each point representing 3-4 pooled assembloids per batch, for 4 batches. Error bars represent mean ± SD. Data were plotted in GraphPad Prism 9.0.0. *p<0.05, **p<0.01, ***p<0.001. **D.** β-galactosidase staining for all the different assembloid conditions. Positive β-galactosidase areas were measured with ImageJ and normalised to the area of the section in each image and batch corrected by normalising to the mean of the values for each batch. Kruskal-Wallis test with Benjamini-Hochberg correction and Dunn’s multiple comparison test was performed. For all conditions n = 6 with each point representing one section per assembloid, per batch, for 3 batches. *p<0.05, **p<0.01, ***p<0.001. Batch correction was applied by normalising each value to the mean of the values for each batch. Outlier removal was performed based on the Inter-Quartile Range (IQR) proximity rule. Data were plotted in R 4.2.2.

Confirming the transcriptomic changes, we were also able to observe significant reduction of the LAMINB1 protein levels, another aging-related phenotype, in the Progerin-overexpressing assembloids (Figure 6.C). Finally, the β-galactosidase staining on assembloid sections, revealed significant higher levels of senescent cells in the Progerin-overexpressing assembloids (Figure 6.D). Overall, we consider that these results reveal a successful induction of aging in the assembloid model with the inducible expression of Progerin.

### Neurodegeneration phenotypes in the Progerin-inducible aged assembloids

Next, we investigated whether the Progerin-overexpressing aged assembloids, at D60 of culture, lead to PD relevant neurodegeneration phenotypes. First, we measured the catecholamine levels in the StrO tissue in assembloids from the control (WT_Untreated, WT_DOX and Progerin_Untreated) and aged conditions (Progerin_DOX) (Figure 7.A). Catecholamine levels were significantly lower in the striatum of the aged assembloid model. These data along with the RNA sequencing analysis, showing the clustering of all the control samples (Figure 6.A), demonstrates a similar phenotype between the 3 control conditions. Therefore, in the following experiments we only compared assembloids generated from the Progerin line without doxycycline treatment (Progerin_Untreated) with assembloids from the same line with doxycycline supplementation for inducing the Progerin overexpression (Progerin_DOX). KEGG and GO enrichment analysis from the bulk RNA sequencing transcriptomic data revealed significant dysregulation of synaptic and DA transmission related pathways (Figure 7.B), with the majority of the responsible genes to be downregulated in the Progerin-overexpressing assembloids (Supplementary Figure 14.A-E). To confirm these results at the protein level, we performed Western blotting for the postsynaptic protein Gephyrin and the presynaptic proteins VAMP2 and Synaptotagmin1 (SYN). Importantly, the three of them were significantly downregulated in Progerin-overexpressing assembloids (Figure 7.C-E).

**Figure 7:**
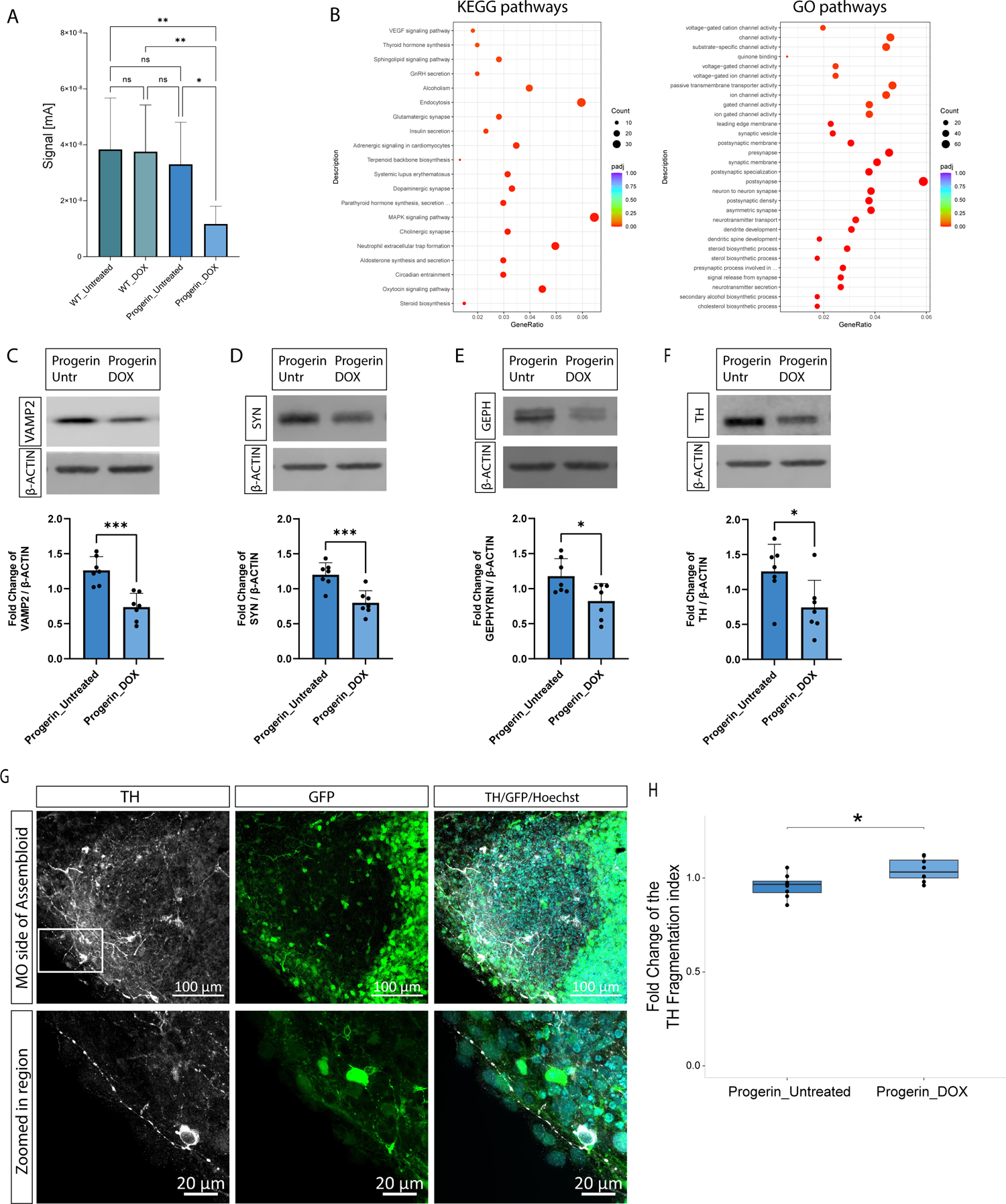
Early neurodegeneration phenotypes in Progerin-overexpressing assembloids. **A.** Bar blot showing the electrochemical measurements in the StrO side of assembloids from the different conditions at D60. One-way ANOVA with Tukey’s multiple comparison test was performed. For all conditions n = the mean of measurements from 5 different positions in 3-4 assembloids per batch for 3 batches (WT_Untreated n = 11, WT_DOX n = 10, Progerin_Untreated n = 11, Progerin_DOX n = 9). Error bars represent mean ± SD. **B.** KEGG and GO pathway enrichment analysis of the DEGs between PROG-DOX and PROG_UNTR samples. **C.** Western blot for the protein levels of VAMP2 normalised to β-Actin. **D.** Western blot for the protein levels of Synaptotagmin1 (SYN) normalised to β-Actin. **E.** Western blot for the protein levels of Gephyrin normalised to β-Actin. **F.** Western blot for the protein levels of TH normalised to β-Actin. **G.** Representative confocal image of the MO side of a 70 μm Progerin-overexpressing assembloid section with TH and Hoechst immunostaining. The white square indicates the zoomed in region showing a representative image of a fragmented TH+ neurite. **H.** Plot showing the TH fragmentation index as quantified by our neuronal skeleton quantification approach with MATLAB (Bolognin et al., 2019). Welch’s t-test was performed with n = 8 for each condition, where each point represents the average of 3-5 sections per assembloid per batch, for 4 batches. In all plots batch correction was applied by normalising each value to the mean of the values for each batch. *p<0.05, **p<0.01, ***p<0.001. For C-F plots, Welch’s t-test was performed in each plot with n = 7 for each condition, where each point represents 3-4 pooled assembloid per batch, for 7 batches. Outliers were calculated in GraphPad Prism 9.0.0 using the ROUT method Q 1%. Error bars represent mean ± SD. For plot H, data were plotted in R 4.2.2 and outlier removal was performed based on the Inter-Quartile Range (IQR) proximity rule.

Loss of TH positive dopaminergic neurons which leads to striatal DA depletion is the key characteristic of PD (Chung et al., 2020). The lower catecholamine levels detected in the striatal compartment of aged assembloids, is a first indication that this pathology hallmark is recapitulated in the aging model. To further substantiate this finding, we assessed the levels of TH protein in the Progerin-overexpressing assembloids via Western blotting. Consistent with the electrochemical measurements, also the TH protein levels are reduced upon induced aging (Figure 7.F). Finally, we investigated the integrity of the dopaminergic neuronal population within the assembloid model. Strikingly, this revealed an increase in the fragmentation of neurites from TH positive dopaminergic neurons, which is an early sign of degeneration (Figure 7.G-H). The TH fragmentation was calculated using our automated image analysis method (Bolognin et al., 2019). Altogether, these data suggest that there is indeed an early loss of dopaminergic neurons’ function in induced aged assembloids that mimics early stages of PD pathology. Accordingly, this new model could be very valuable to investigate early alterations during the onset and progression of PD.

## Discussion

In this study, we developed a midbrain-striatum assembloid model that resembles the physiological nigrostriatal pathway connectivity. This model represents a relevant tool for the investigation of PD. To date, most studies on PD research with advanced cellular 3D models are based on the use of midbrain organoids (Becerra-Calixto et al., 2023; H. Kim et al., 2019; Monzel et al., 2017; Smits et al., 2019). Although these organoids have proven to be an excellent tool for studying the dopaminergic neurons’ vulnerability in different PD-related conditions, they are not able to recapitulate the dysregulated connectivity in the nigrostriatal pathway, which is crucially affected in PD (Fuxe et al., 2006; Singh et al., 2015). For the reconstruction of the nigrostriatal pathway connectivity, here we show the generation of a midbrain-striatum assembloid model that retains the identity of the two regions with the expression of midbrain and striatum specific markers respectively.

Single nuclei RNA sequencing analysis further revealed the identity specificity of MOs and StrOs, while the assembloid model demonstrated the preservation of cellular populations from both organoids, with additionally the identification of A9 and A10 dopaminergic neuronal clusters that are not clearly detected in our MOs dataset. A9 dopaminergic neurons were revealed by the high expression of *KCNJ6* that encodes GIRK2 protein (H. Li et al., 2022), while expression of *OTX2* was found only in the A10 cluster. *OTX2* encodes a transcription factor that is crucial for the specification of the A10 dopaminergic neurons (Grealish et al., 2014). In parallel, the MSN population in the assembloid model is specified by the co-expression of *ARPP21*and *PPP1R1B*, a strong indication of an MSN identity (Ivkovic & Ehrlich, 1999; Straccia et al., 2015), which is not evident in the StrOs dataset where only *ARPP21* expression was found. This illustrates that the assembloid model, probably through a functional communication between both regions, favours the further specification of the midbrain dopaminergic neurons as well as the MSNs of the striatum.

Enrichment analysis of the DEGs between assembloids and MOs, and assembloids and StrOs, revealed the upregulation of genes related to developmental neurogenesis, axonal guidance, and neuronal maturity. In line with the importance of brain’s interregional communication in neuronal maturity and functionality (Qin et al., 2015; Voytek & Knight, 2015), it is possible that neurogenesis and neuronal maturity are promoted by the interactions formed between midbrain and striatal neurons in the assembloid model. These observations align with another study in cortico-thalamic assembloids, where neuronal maturity was observed in the assembloid level compared to individual organoids (Xiang et al., 2019). Moreover, this assumption is further supported by the reduced expression of cellular and oxidative stress related genes in midbrain-striatum assembloids, as elevated stress levels in cerebral organoids have been linked with impaired cellular maturity (Bhaduri et al., 2020).

Nigrostriatal pathway connectivity and functionality are also confirmed in assembloids, with the formation of active synapses between midbrain and striatal neurons and the release of catecholamines from the midbrain to striatum. This is specifically important, as the catecholamine DA release in the dorsal striatum from the SNpc dopaminergic neuronal axons is the major functionality of the nigrostriatal pathway and it is crucial for behaviour and movement control (Aarts et al., 2011; Sulzer et al., 2016).

A limitation of iPSC-derived organoid models in the research of age-related neurodegenerative diseases is that their developmental nature lacks the aging-specific phenotypes (Simpson et al., 2021). Since aging is the major risk factor in PD, we aimed to model this aspect in midbrain-striatum assembloids. To achieve this, we leveraged the overexpression of Progerin for introducing aging characteristics to our model. Similar approach has been previously described, where transient Progerin overexpression in iPSC derived dopaminergic neurons resulted in aging and neurodegeneration phenotypes (Miller et al., 2013). In our approach for Progerin overexpression, we used iPSCs carrying a Progerin transgene under the control of the Tet-On system (Gabassi et al., *In Preparation*), where doxycycline supplementation is essential for the inducible expression. Our optimised concentration of doxycycline for Progerin overexpression can be safely used in our system, as it does not affect the viability and metabolic activity in assembloids, and it does not introduce major changes in their transcriptome profile. Progerin overexpression was achieved in approximately half of the cells in doxycycline treated assembloids carrying the Progerin transgene, creating a mosaic Progerin-overexpressing model. In this mosaic model, we found that Progerin-overexpressing cells acquire aging characteristics compared to the non-Progerin-expressing cells. Some of these characteristics are the increased levels of DNA double strand breaks marked with the double positive H2AX-53BP1 nucleus foci (Shibata & Jeggo, 2020). Moreover, Progerin-overexpressing cells and neurons had significantly higher levels of p21^CIP1^, p53 and p16^INK4A^. The upregulation of these three proteins has been shown to be associated with cellular senescence and aging (Kumari & Jat, 2021; Wagner & Wagner, 2022).

To assess how Progerin-overexpressing cells affect the whole assembloid model we performed transcriptomic analysis with bulk RNA sequencing. Clustering of the data based on their transcriptomic similarity and the differential gene expression revealed a distinct transcriptomic profile for assembloids with Progerin overexpression. To investigate whether there are aging-related transcriptomic differences we performed a benchmarking analysis with the use of human brain post-mortem transcriptomic data from aged individuals (Berchtold et al., 2008; González-Velasco et al., 2020). An aging-related transcriptomic profile was observed in Progerin-overexpressing assembloids, with the identification of aging-related genes with a common expression pattern between assembloids and human brain post-mortem data. Noteworthy, Progerin assembloids DEG showed *PNOC* downregulation and *GMPR* upregulation, that were found to correlate best with the biological age of the human brain post-mortem data (Velasco et al., 2020).

Significantly upregulated genes found in Progerin assembloids have also been associated with aging and neurodegeneration. Specifically, higher expression of *CHI3L1* has been linked to aging and neurodegeneration (Moreno-Rodriguez et al., 2020; Sanfilippo et al., 2019), *COL21A1* upregulation has been correlated with Alzheimer’s disease (AD) (Kong et al., 2009) and *IER3* gene has been implicated in cellular stress response and it is upregulated in inflammatory conditions (Arlt & Schäfer, 2011). From the significant downregulated genes*, PNOC* and *VIP* are associated with inhibitory neurotransmission in GABAergic neurons and are both significantly downregulated in human aged brains (Loerch et al., 2008). *SST* and *CRH* downregulation has been found in aging brain but also in AD genetic signature (Berchtold et al., 2013; Loerch et al., 2008; Peng et al., 2021). Similarly, downregulation of *NRGN* expression correlates with higher density of amyloid plaques in post mortem brains of AD patients (Sun et al., 2021). *GRM4* which encodes for the Glutamate Metabotropic Receptor 4 has also been found to be reduced in the prelimbic of aged rats (Hernandez et al., 2018). *KCNAB1* and *GABRA4* have a positive co-expression with *BDNF* which shows gradual downregulation in aging human prefrontal cortex (Oh et al., 2016). Additionally, *DRD1* and *DRD2* DA receptors’ downregulation correlates with brain aging (Lubec et al., 2021). Both receptors are crucial for DA signalling in the striatum, and their reduced expression can be linked to motor and cognitive abnormalities (Hemby et al., 2003). *DLX6* encodes an important transcription factor for the regulation of GABAergic neurons (Lombares et al., 2019) and therefore its downregulation could be associated with aging. *NETO1* is essential for the regulation of the connectivity of glutamatergic neurons (Orav et al., 2017; Straub et al., 2011) and its downregulation could cause dysfunction in synaptic circuits. Reduction of *COBL* has be linked with reduced dendrite arborisation (Ahuja et al., 2007). Lastly, downregulation of *CHCHD2* could be associated with the aging brain and development of PD, as studies have shown that mutations in this gene are tied to dysfunctional mitochondria (Kee et al., 2021; Meng et al., 2017), while at the same time reduction of *CHCHD2* mRNA levels has been found in erythrocytes of PD patients (Liu et al., 2021). The aging phenotype observed in the transcriptomic profile of the Progerin-overexpressing assembloids was further validated by the significant reduction of the LAMINB1 protein levels and the significant increase in senescence-activated β-galactosidase positive areas. Both these phenotypes have been considered hallmarks of aging and senescence (Dodig et al., 2019; Matias et al., 2022; Miller et al., 2013).

Many of the genes mentioned previously are not only related to aging but also to neurodegenerative diseases such as AD and PD, highlighting the interconnected relationship between aging and neurodegeneration. Synaptic dysfunction in the aging brain is one of the major promoters of neurodegeneration (Azam et al., 2021; Buss et al., 2021; Talyansky & Brinkman, 2021). Dysregulation of synaptic and DA neurotransmission pathways revealed in the transcriptomic data of Progerin-overexpressing assembloids, was further validated by the downregulation of the post-synaptic protein Gephyrin, an important scaffolding protein that regulates the organisation of post-synapses in striatal GABAergic neurons (Choii & Ko, 2015), and the pre-synaptic proteins VAMP2 and Synaptotagmin1. VAMP2 regulates the fusion of synaptic vesicles and neurotransmitter release through the SNARE complex (Yan et al., 2022) and Synaptotagmin1 is an essential Ca^2+^ sensor for the fast release of DA (Banerjee et al., 2020, 2022). In parallel, catecholamine measurements showed significantly lower catecholamine levels in the striatum of the Progerin-overexpressing assembloids, indicating dysregulation of DA release in the aged-induced model, a PD-relevant phenotype (Jiang et al., 2019). These results reveal a defective synaptic and DA release system in Progerin-overexpressing assembloids.

Moreover, low TH protein levels in Progerin-overexpressing assembloids could be linked to dysregulated DA synthesis system and the presence of vulnerable dopaminergic neurons. TH is the key enzyme of DA synthesis (Daubner et al., 2011). Lower TH protein levels have been previously observed in surviving dopaminergic neurons of PD patients, indicating their vulnerability to the disease (Kastner et al., 1993). Moreover, loss of TH activity followed by TH protein decline has been linked to DA deficiency in PD (Tabrez et al., 2012). Additionally, quantification of the fragmentation index of the TH+ neurites revealed that dopaminergic neurons in the age-induced assembloids are more fragmented, demonstrating a degeneration phenotype prior to their neuronal death (Lin et al., 2016). The axonal fragmented phenotype has been previously described in iPSC-derived PD neuronal models (Kouroupi et al., 2017). These results support that our age-induced assembloids display an early PD-relevant neurodegeneration phenotype, with dysregulation in the DA synthesis and release systems indicative of axonal degeneration preceding the eventual loss of dopaminergic neurons’ soma, in retrograde fashion as typically observed in PD (Cheng et al., 2011; Chu et al., 2012; L. H. Li et al., 2009; Miwa et al., 2005).

The here presented age-induced midbrain-striatum assembloid model offers new possibilities for PD research. Given the crucial role of aging in PD development, it is imperative to diligently consider its influence when studying the disease’s phenotypes through *in vitro* models. By incorporating aging characteristics in cellular models derived from PD patient cell lines, we can more accurately recapitulate the disease state of patients and unveil cellular dysfunctionalities that remain unnoticed in the current developmental models. Studies with cell lines sourced from PD patients in the future will provide further insights into the model’s robustness and its potential applications in personalised medicine.

## Authors Contribution

K.B. designed and executed experiments, analysed and interpreted data, prepared figures, and wrote the original draft. C.S. designed and executed experiments for the striatum organoids optimisation, reviewed and edited the manuscript. G.G.G. contributed on the optimisation of the Rabies monosynaptic tracing experiment in assembloids. E.G. contributed to the generation and quality control of the Progerin iPSCs. S.S. and K.B. performed the electrochemical measurements in organoids and assembloids. J.E.R.G. performed the whole mount imaging of assembloids and analysis. P.A. supervised the high-content imaging workflow. G.R. contributed to the analysis of microelectrode array data. F.P. generated the RBV and LV viruses for the Rabies monosynaptic tracing experiment. U.K., P.E. and R.M. reviewed and edited the manuscript. A.S. and F.E. provided scientific feedback in regular project meetings, reviewed, and edited the manuscript. J.C.S. conceived and supervised the project and edited the manuscript.

## Acknowledgements

This study was supported by the Luxembourg National Fund FNR-Inter and the Austrian Science Fund FWF (FWF-INTER – INTER/FWF/19/14117540/PDage; FWF-SFB F78/P1040-016-015).

Special thanks to Prof. Alexander Skupin for his scientific input on single nuclei RNA sequencing analysis data and Dr. Henry Kurniawan for his feedback on the flow cytometry experiment. Rights retention statement: This research was funded in whole by the FNR-Luxembourg. For the purpose of Open Access, the author has applied a CC BY public copyright license to any Author Accepted Manuscript (AAM) version arising from this submission.

## Ethical approval

Ethics Review Panel (ERP) of the University of Luxembourg and the national Luxembourgish research ethics committee (CNER, Comité National d’Ethique de Recherche) have approved the work with induced pluripotent stem cells (iPSCs). CNER No. 201901/01; ivPD

