## Supplementary Information for "Age-induced midbrain-striatum assembloids model early phenotypes of Parkinson’s disease"

**Supplementary Figure Legends**

**Supplementary Figure 1: Characterisation of iPSC and NESC lines with immunofluorescence staining.** iPSCs and NESCs were immunostained for the detection of pluripotent and neuroepithelial stem cell markers respectively. **A.** iPSCs of the 200 cell line were immunostained for SSEA-4, NANOG, OCT4 and Hoechst. **B.** iPSCs of the 201 cell line were immunostained for SSEA-4, OCT4, NANOG, TRA1-60 and Hoechst. **C.** iPSCs of the 389 and 390 cell lines were immunostained for SSEA-4, OCT4 and Hoechst. **D.** iPSCs of the 389 and 390 cell lines were immunostained for NANOG, TRA1-60 and Hoechst. **E.** NESCs of the 201 and 200 cell lines were immunostained for PAX6, NESTIN and Hoechst. These lines were previously genetically engineered for the intrinsic expression of GFP. **F.** NESCs of the 201 and 200 cell lines were immunostained for SOX1, SOX2 and Hoechst. These lines were previously genetically engineered for the intrinsic expression of GFP. **G.** NESCs of the 389 and 390 cell lines were immunostained for PAX6, SOX2, NESTIN and Hoechst.

**Supplementary Figure 2: Different conditions used for the generation of striatum organoids. A-D.** Schematic representation of the different steps in the differentiation process of StrOs generation in the condition C4 (A)**,** in the condition C3 (B), in the condition RA (C) and in the condition SR (D).

**Supplementary Figure 3: Striatum organoids characterisation. A.** qPCR plots showing the relative expression of progenitor markers specific to the LGE and striatum brain regions. For *FOXG1*, *ASCL1*, *GSX2* and *CTIP2*, Kruskal-Wallis test with Benjamini-Hochberg correction and Dunn's multiple comparison test was performed. For *FOXP1* and *FOXP2*, one-way ANOVA, with Tukey's multiple comparison test was performed. **B.** qPCR plots showing the relative expression of progenitor markers specific to the MGE region of the brain. For *NKX2.1*, Kruskal-Wallis test with Benjamini-Hochberg correction and Dunn's multiple comparison test was performed. For *OTX2*, one-way ANOVA, with Tukey's multiple comparison test was performed. **C.** qPCR plots showing the relative expression of striatum specific markers. For *DARPP32*, one-way ANOVA, with Tukey's multiple comparison test was performed. For *DRD1* and *DRD2*, Kruskal-Wallis test with Benjamini-Hochberg correction and Dunn's multiple comparison test was performed. **D.** Western blot showing the protein levels of DARPP32 in the different conditions of StrOs. One-way ANOVA, with Tukey's multiple comparison test was performed. **E.** Western blot showing the protein levels of DRD1 in the different conditions of StrOs. One-way ANOVA, with Tukey's multiple comparison test was performed. **F.** Western blot showing the protein levels of DRD2 in the different conditions of StrOs. One-way ANOVA, with Tukey's multiple comparison test was performed. **G.** Western blot showing the protein levels of GAD65 in the different conditions of StrOs. One-way ANOVA, with Tukey's multiple comparison test was performed. For all plots, n = 3-5, where each point is one sample of pooled 6-8 pooled organoids, from 3-5 batches. *p<0.05, **p<0.01, ***p<0.001. qPCR data were normalised to ACTINB expression and fold change to the C4_D35 condition was calculated. For the western blots, data were normalised to β-Actin levels and batch correction was applied by normalising each value to the mean of the values for each batch. Error bars represent mean ± SD. In all plots, outliers were calculated in GraphPad Prism using the ROUT method Q 1%. Data were plotted in GraphPad Prism 9.0.0.

**Supplementary Figure 4: Co-culture media optimisation for the assembloid model. A.** Schematic representation of the protocol used for the generation of assembloids. **B.** Table showing the different media that were tested for the optimisation of the co-culture condition. **C.** Immunostaining of MO 70 µm sections for the expression of TH, cultured for 20 days in the different co-culture conditions. Images were acquired with the Yokogawa high content imaging microscope. Kruskal-Wallis test with Benjamini-Hochberg correction and Dunn's multiple comparison test was performed. N = 6, where each point represents the average of 2 sections per organoids per batch, for 3 batches. Batch correction was applied by normalising each value to the mean of the values for each batch. $^*p<0.05$, $^{**}p<0.01$, $^{***}p<0.001$, $^{****}p<0.001$. Outlier removal was performed based on the Inter-Quartile Range (IQR) proximity rule. Data were plotted in R 4.2.2.

**Supplementary Figure 5: Immunostaining for TH in assembloids cultured in the different co-culture conditions.** 70 µm assembloid sections cultured for 20 and 35 days in the different co-culture and StrO conditions, were immunostained with TH and Hoechst for the quantification of the TH positive cells. Images were acquired with the Yokogawa high content imaging microscope. Kruskal-Wallis test with Benjamini-Hochberg correction and Dunn's multiple comparison test was performed. N = 6-9, where each point represents one section, per assembloid, per batch, for 3 batches. In all plots batch correction was applied by normalising each value to the mean of the values for each batch. $^*p<0.05$, $^{**}p<0.01$, $^{***}p<0.001$. Outlier removal was performed based on the Inter-Quartile Range (IQR) proximity rule. Data were plotted in R 4.2.2.

**Supplementary Figure 6: Immunostaining for DARPP32 in assembloids cultured in the different co-culture conditions.** 70 µm assembloid sections cultured for 20 and 35 days in the different co-culture and StrO conditions, were immunostained with DARPP32 and Hoechst for the quantification of the DARPP32 positive cells. Images were acquired with the Yokogawa high content imaging microscope. Kruskal-Wallis test with Benjamini-Hochberg correction and Dunn's multiple comparison test was performed. N = 6-9, where each point represents one section, per assembloid, per batch, for 3 batches. In all plots batch correction was applied by normalising each value to the mean of the values for each batch. $^*p<0.05$, $^{**}p<0.01$, $^{***}p<0.001$. Outlier removal was performed based on the Inter-Quartile Range (IQR) proximity rule. Data were plotted in R 4.2.2.

**Supplementary Figure 7: Markers expression in the different clusters of assembloids, identified by single nuclei RNA sequencing analysis. A-I.** Dot plots showing the expression of markers specific to dopaminergic neurons (DANs) (A-B), mature neurons (C), GABAergic neurons (D), medium spiny neurons (MSNs) (E-F), LGE progenitors (G), radial glia cells (RGCs) (H) and glia progenitors (I).

**Supplementary Figure 8: Single nuclei RNA sequencing analysis of midbrain organoids. A.** UMAP embedding of the different cellular clusters in MOs. **B.** Percentage of the cellular composition in MOs. **C**. Unsupervised hierarchical clustering of cell clusters, using the average expression with Z-score normalisation of the top 500 most variable genes. **D.** Spearman's correlation between the different cell types in MOs. **E-J.** Dot plots showing the expression of markers specific to dopaminergic neurons (E-F), GABAergic neurons (G), mature neurons (H), oligodendrocyte precursor cells (OPCs) (I) and radial glia cells (RGCs) (J).

**Supplementary Figure 9: Single nuclei RNA sequencing analysis of striatum organoids. A.** UMAP embedding of the different cellular clusters in StrOs. **B.** Percentage of the cellular composition in StrOs. **C**. Unsupervised hierarchical clustering of cell clusters, using the average expression with Z-score normalisation of the top 500 most variable genes. **D.** Spearman's correlation between the different cell types in StrOs. **E-K.** Dot plots showing the expression of markers specific to mature neurons (E), Telencephalic progenitors (F), MSNs progenitors (G), dopaminergic neurons (H), GABAergic neurons and MSNs (I-J) and GABAergic interneurons (K).

**Supplementary Figure 10: Single nuclei RNA sequencing integration analysis. A.** UMAP embedding of the integrated object, showing the predefined cellular identities in each model. **B.** Spearman's correlation between the different cell types. **C**. Expression of genes related to cellular and oxidative stress response in assembloids compared to MOs and StrOs. Dots represent single cells. Two-sided Wilcoxon test was performed between assembloids-MOs and assembloids-StrOs. $^*p<0.05$, $^{**}p<0.01$, $^{***}p<0.001$, $^{****}p<0.0001$. Data were plotted in R 4.2.2.

**Supplementary Figure 11: Nigrostriatal connectivity in the assembloid model. A.** Representative confocal images of MO and StrO 70 μm sections immunostained for TH, MAP2 and Hoechst. **B.** Representative light sheet microscopic image (12X objective) of a whole assembloid and a ROI. The assembloid was immunostained with Hoechst and TH. The ROI shows the GFP+/TH+ neuron's soma in the MO-GFP side of the assembloid with TH+ projections towards the striatum side in the different planes (Z.236-280) along the Z stack. White arrowheads show the progression of the TH+ projection in images from the different planes. **C.** Representative light sheet microscopic image of a whole assembloid (4X objective) and the ROI (25X objective), in light sheet fluorescence microscopy. The assembloid was immunostained with Hoechst and TH. The ROI shows the GFP+/TH+ neuron's soma in the MO-GFP side of the assembloid with TH+ projections towards the striatum side in the different planes along the Z stack (Z.660-755). White arrowheads show the progression of the TH+ projection in images from the different planes. 3D reconstruction of the neuron across the planes shows the complete TH+ neuronal projection. **D.** Schematic representation of the electrochemical measurement set up in assembloids. **E.** Barplot showing the electrochemical measurements of catecholamines in tissue approximate to the MO and StrO sides of the assembloid model. Welch's t-test was performed. N = 8, where each point represents the average of the 3 technical electrochemical measurements in the MO and StrO side respectively, in each assembloid, for 2 batches. Background measurement was not subtracted in these measurements. Data were plotted in GraphPad Prism 9.0.0. *p<0.05, **p<0.01, ***p<0.001. Error bars represent mean ± SD. **F.** Bar plot showing the electrochemical measurements of catecholamines in StrOs cultured in their normal conditions and StrOs cultured in pre-used media in MOs culture. Welch's t-test was performed. N = 3-8, where each point represents the average measurement value with the background subtracted (PBS measurements) of each organoid per batch, for 3 batches. Data were plotted in GraphPad Prism 9.0.0. *p<0.05, **p<0.01, ***p<0.001. Error bars represent mean ± SD. **G.** Representative confocal images of a 70 μm assembloid section showing the GFP and RFP positive cells from the LV-GP-TVA-GFP and RBV-ΔG-EnvA-RFP infections respectively and immunostained with Hoechst and TH. Close up images indicated by the white square, show the colocalization of Rabies and TH signal in the MO side of the assembloid.

**Supplementary Figure 12: Toxicity assessment of doxycycline. A.** Boxplot showing the quantification of the LDH luminescence signal normalised to the average area of assembloids. One-way ANOVA, with Tukey's multiple comparison test was performed. N = 3, where each point represents the average of 3 technical replicates per batch, for 3 batches. Statistics show the pairwise comparisons of interest between the treatments in the same condition (Progerin or WT) and between the conditions with the same treatment. **B.** Boxplot showing the quantification of the ATP luminescence signal normalised to the average area of assembloids. One-way ANOVA, with Tukey's multiple comparison test was performed. N = 3, where each point represents the average of two technical replicates per batch, for 3 batches. *p<0.05, **p<0.01, ***p<0.001. Data were plotted in R 4.2.2.

**Supplementary Figure 13: Differential expression gene profile in aged and young assembloids. A.** Heatmap showing the hierarchical clustering of samples with similar expression patterns, after Z-score homogenisation of the expression data. **B.** Histogram showing the number of differential genes (up-regulated and down-regulated) for each comparison.

**Supplementary Figure 14: Expression of genes relevant to the synaptic pathways in GO and KEGG enrichment analysis.** Fold change expression of genes in Progerin-overexpressing assembloids (PROG_DOX) versus non-Progerin-expressing assembloids (PROG_UNTR), responsible for the enrichment of Dopaminergic Synapse KEGG pathway (A), Neuron to neuron synapses GO pathway (B), Neurotransmitter secretion GO pathway (C), Postsynapses GO pathway (D) and Presynapses GO pathway (E).

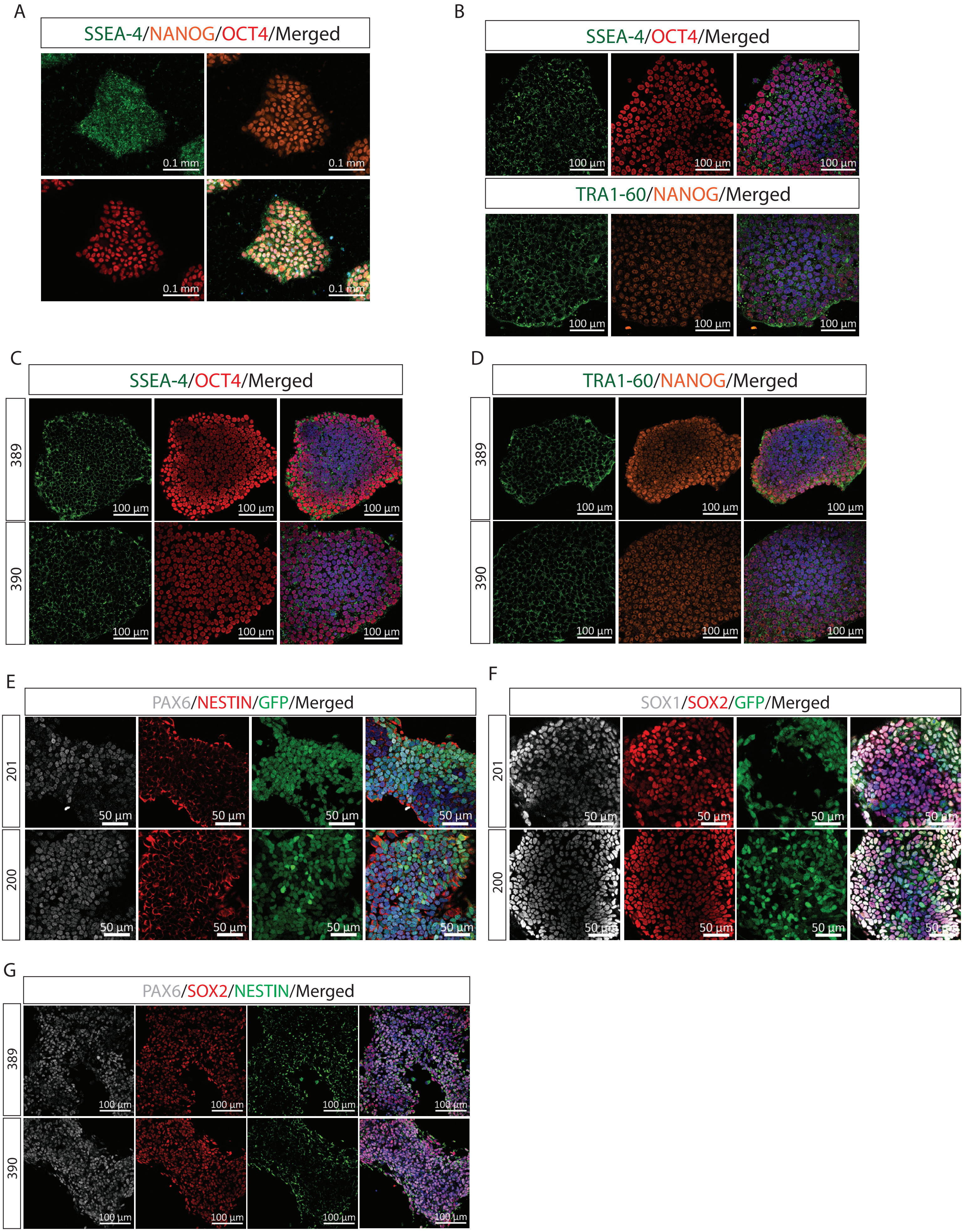


Supplementary Figure 1

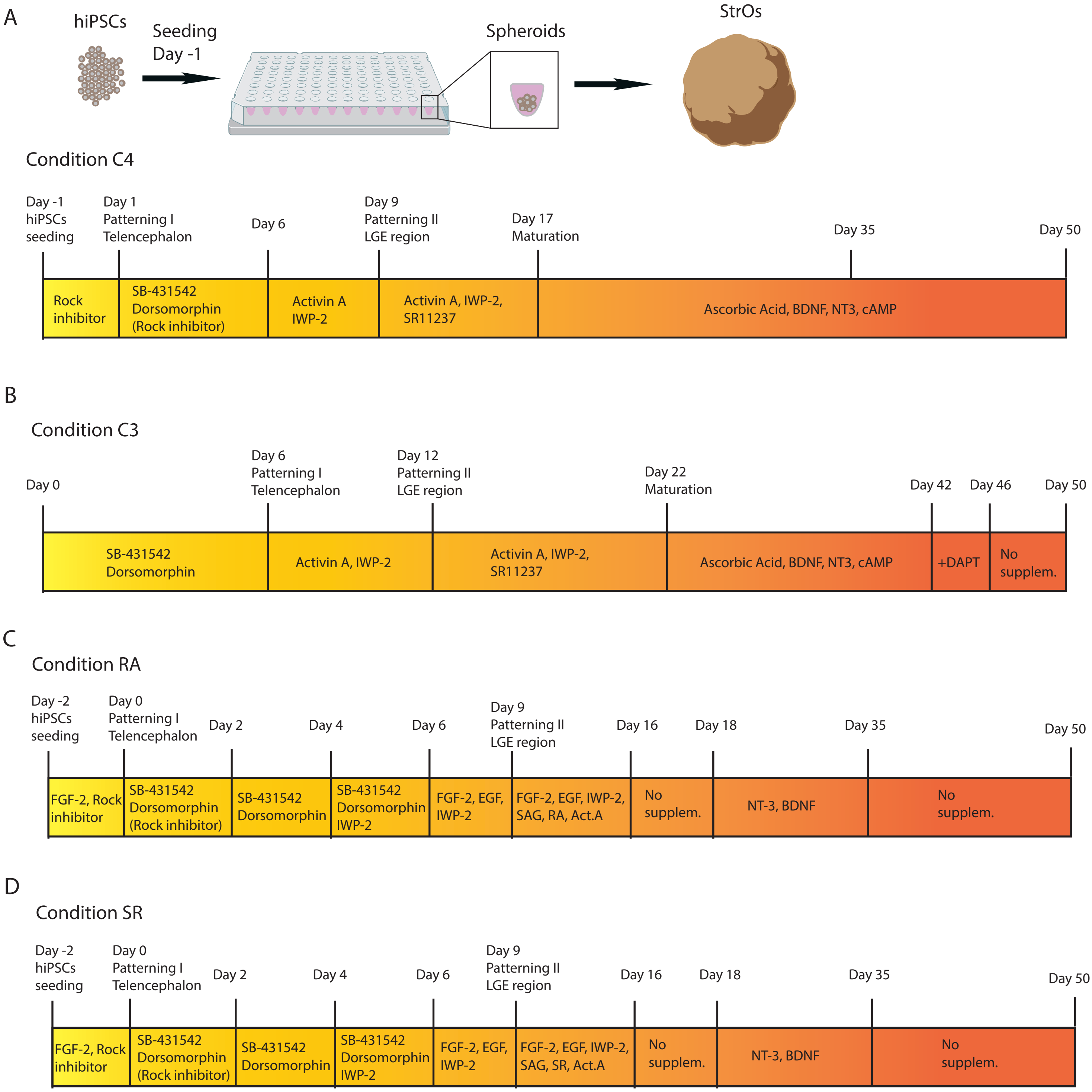


Supplementary Figure 2

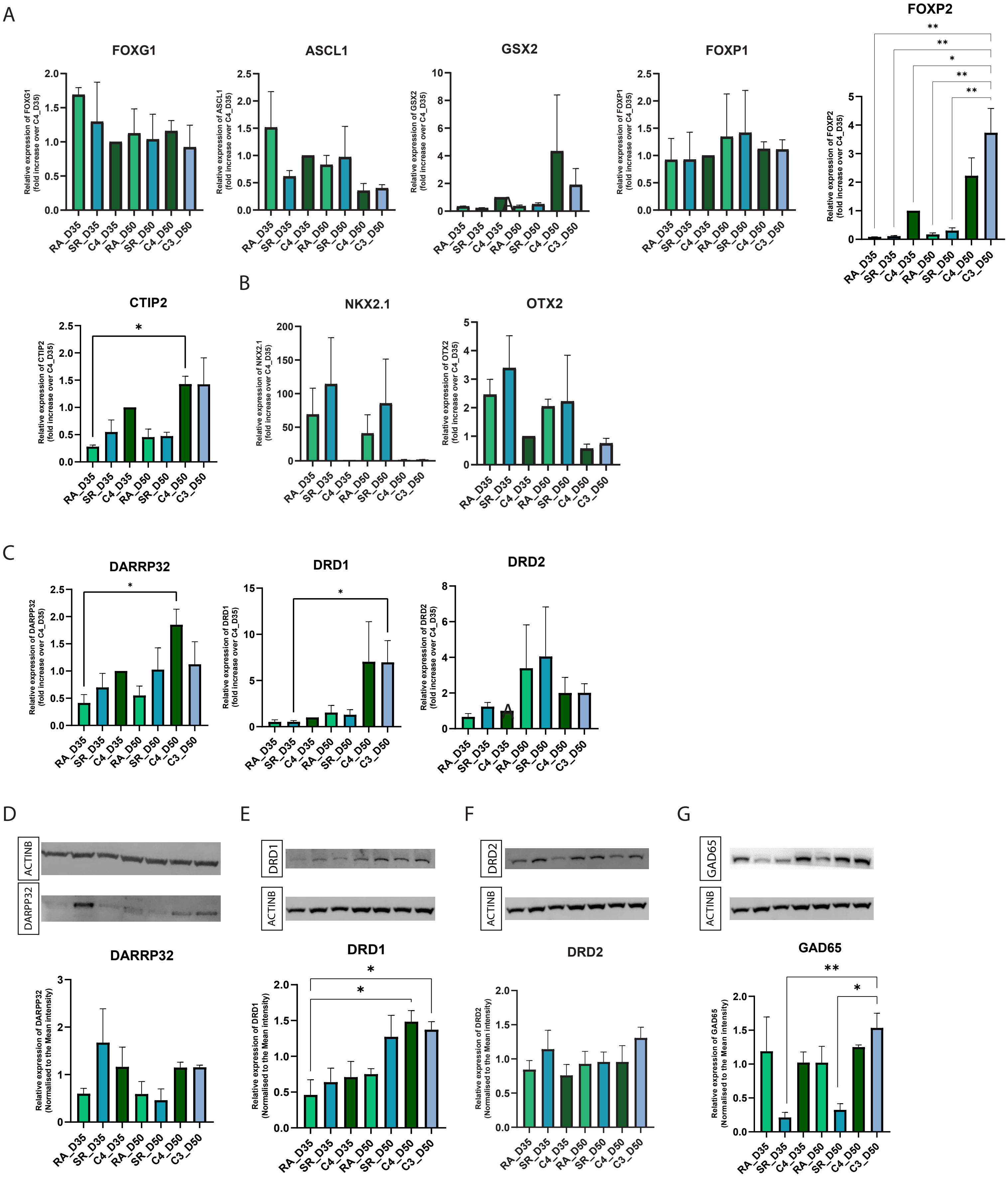


Supplementary Figure 3

A

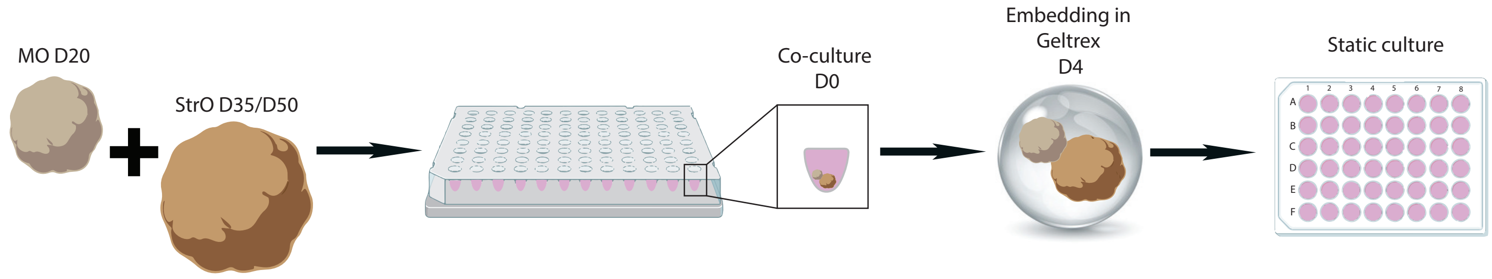


B

| | Media Composition | Small Molecules |
|---|---|---|
| Neural Medium | Neurobasal A, B27, Glutamax, P/S | - |
| Neural Medium Plus | Neurobasal A, **B27 plus**, Glutamax, P/S | - |
| Neural Medium ++ | Neurobasal A, B27, Glutamax, P/S | AA, db cAMP, BDNF, GDNF,NT3 |
| N2B27 ++ | Neurobasal, DMEM/F12, B27, N2, Glutamax, P/S | AA, db cAMP, BDNF, GDNF,NT3 |

C

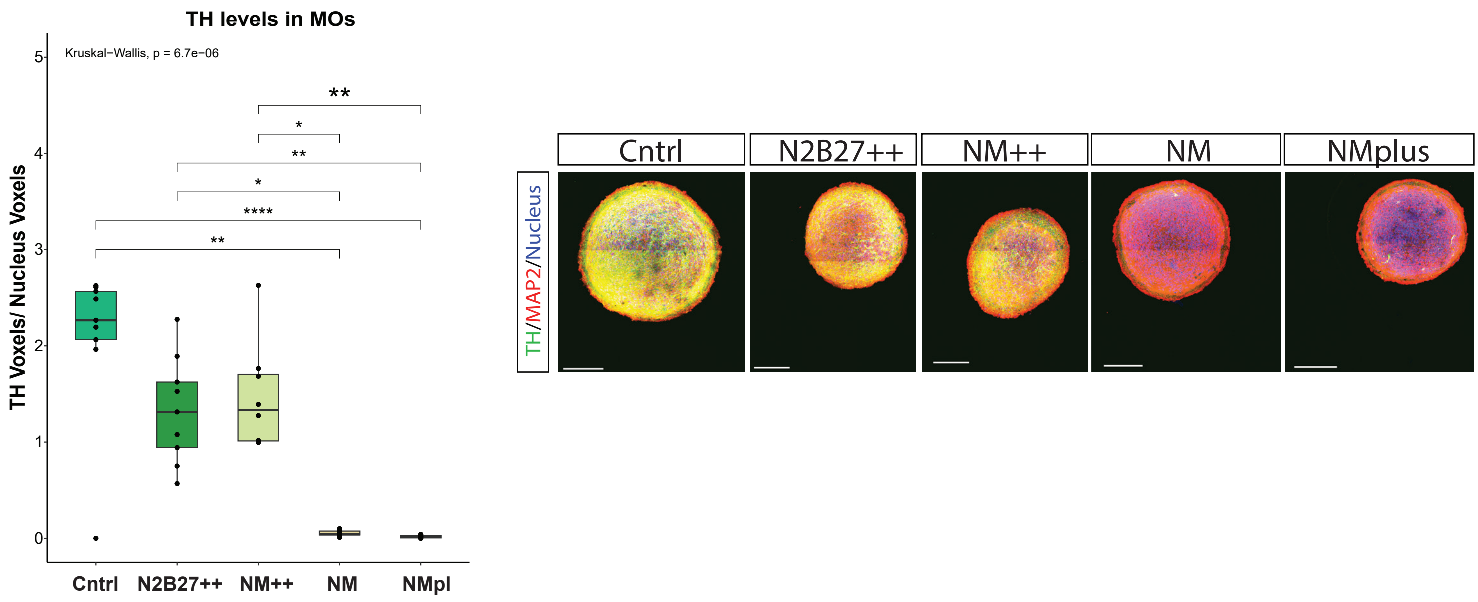


Supplementary Figure 4

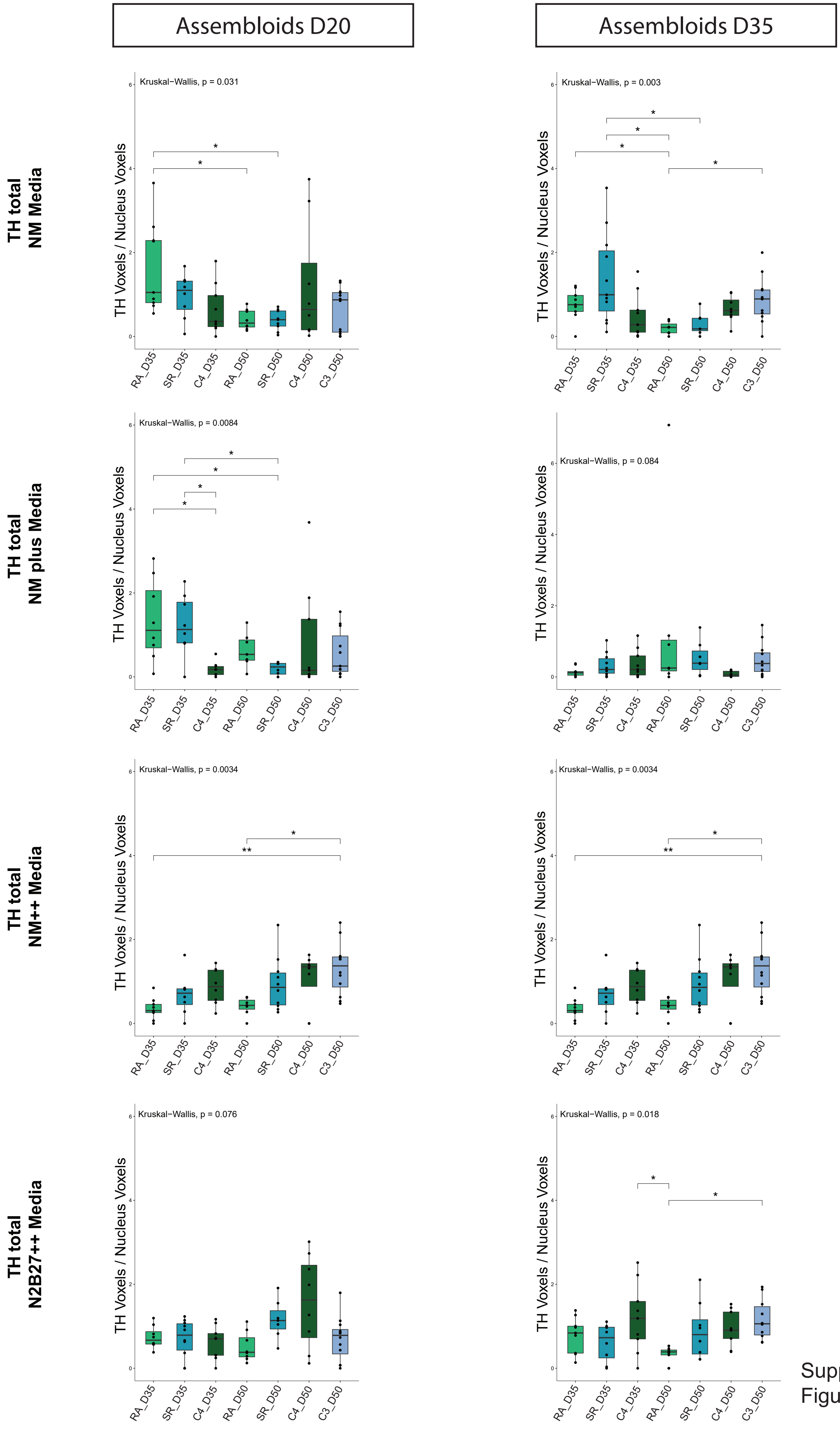


Supplementary Figure 5

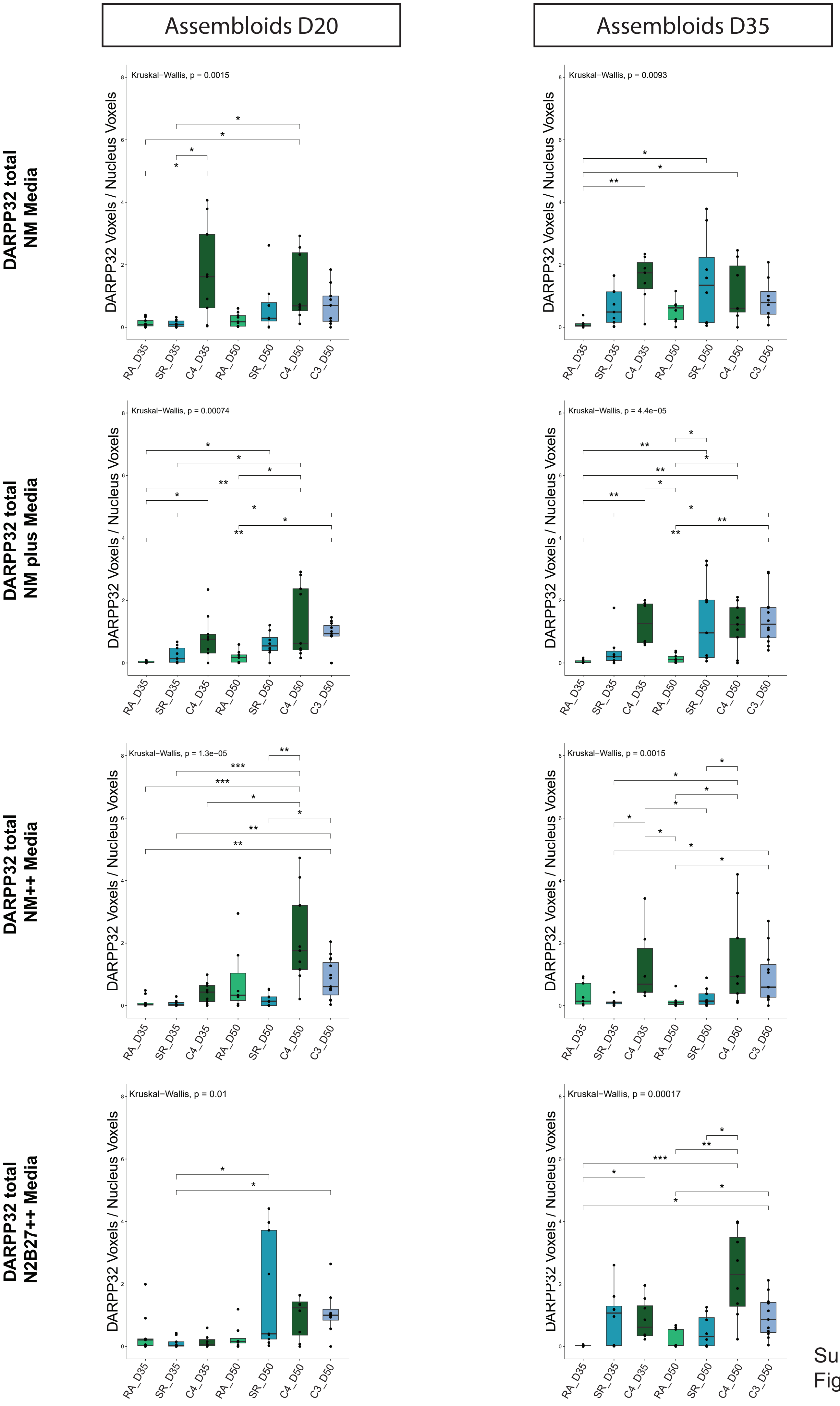


Supplementary Figure 6

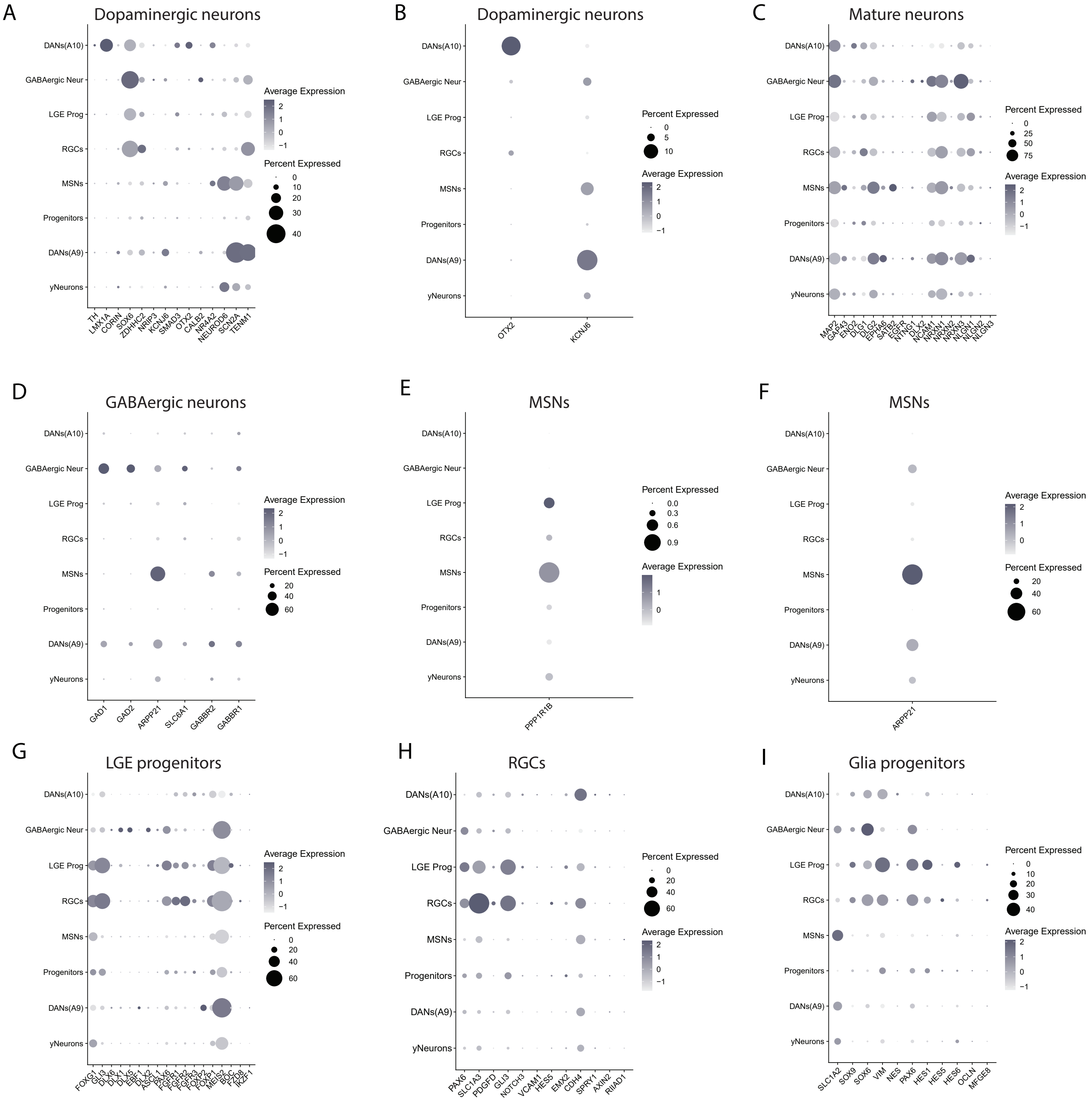


Supplementary Figure 7

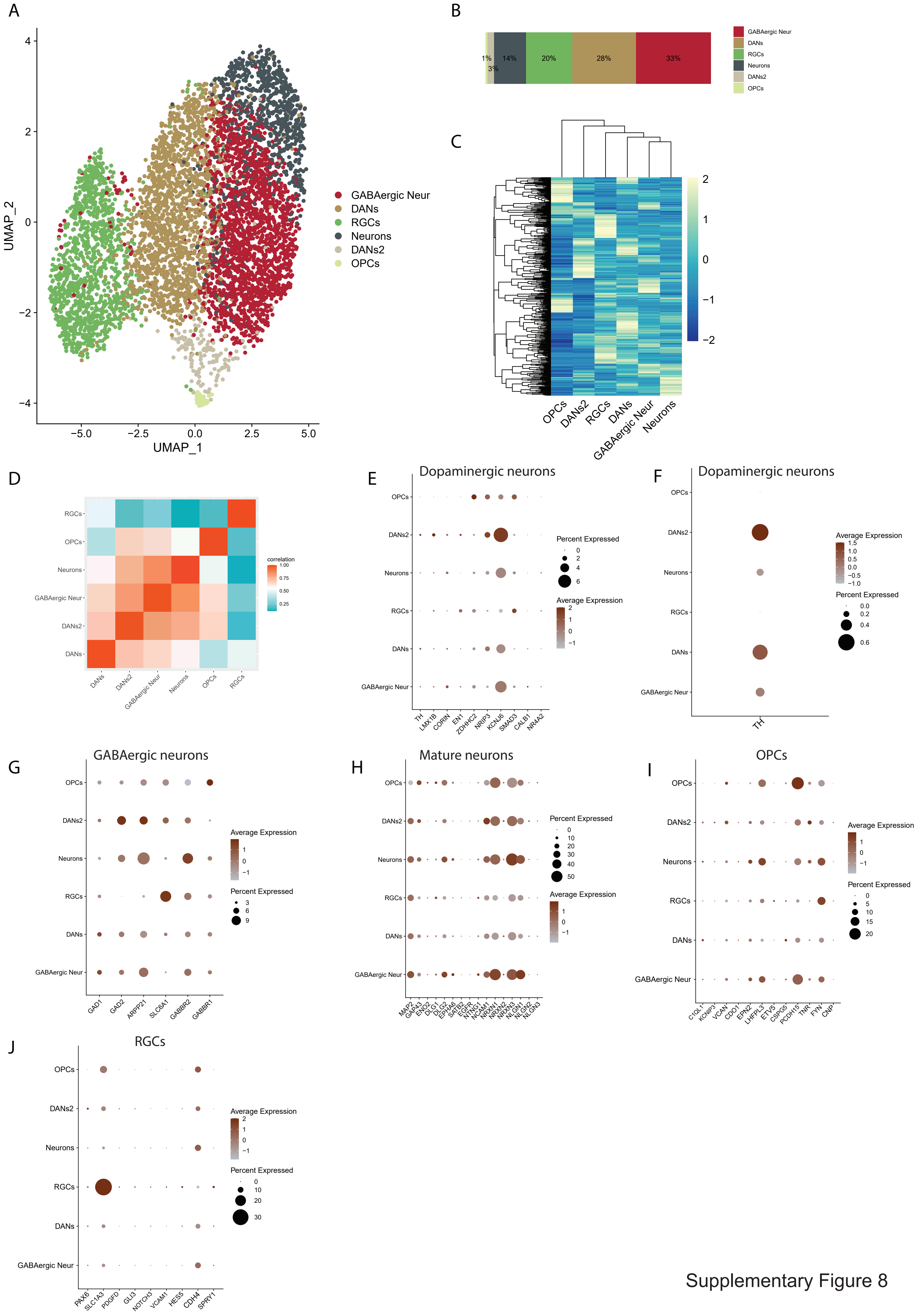


Supplementary Figure 8

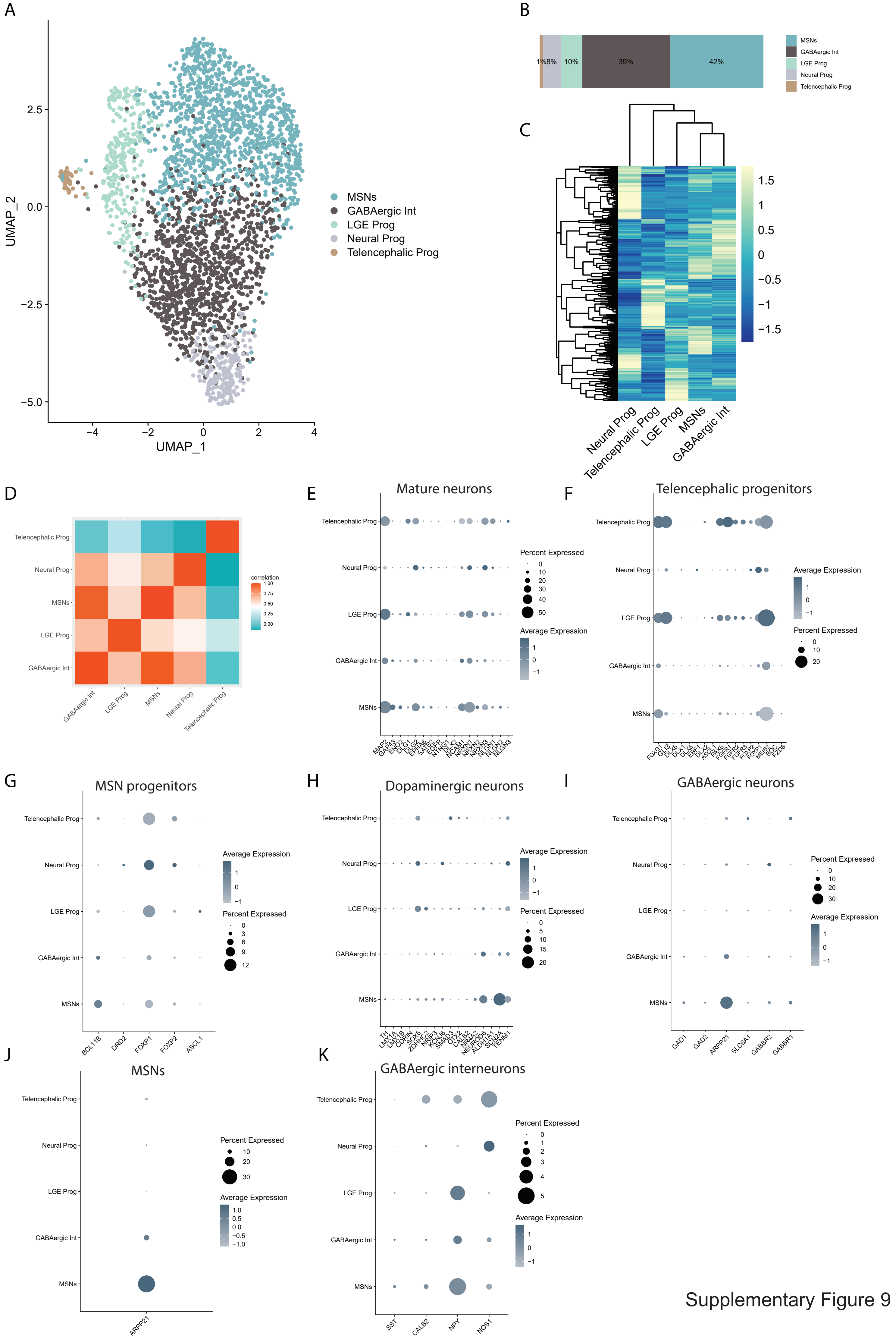


Supplementary Figure 9

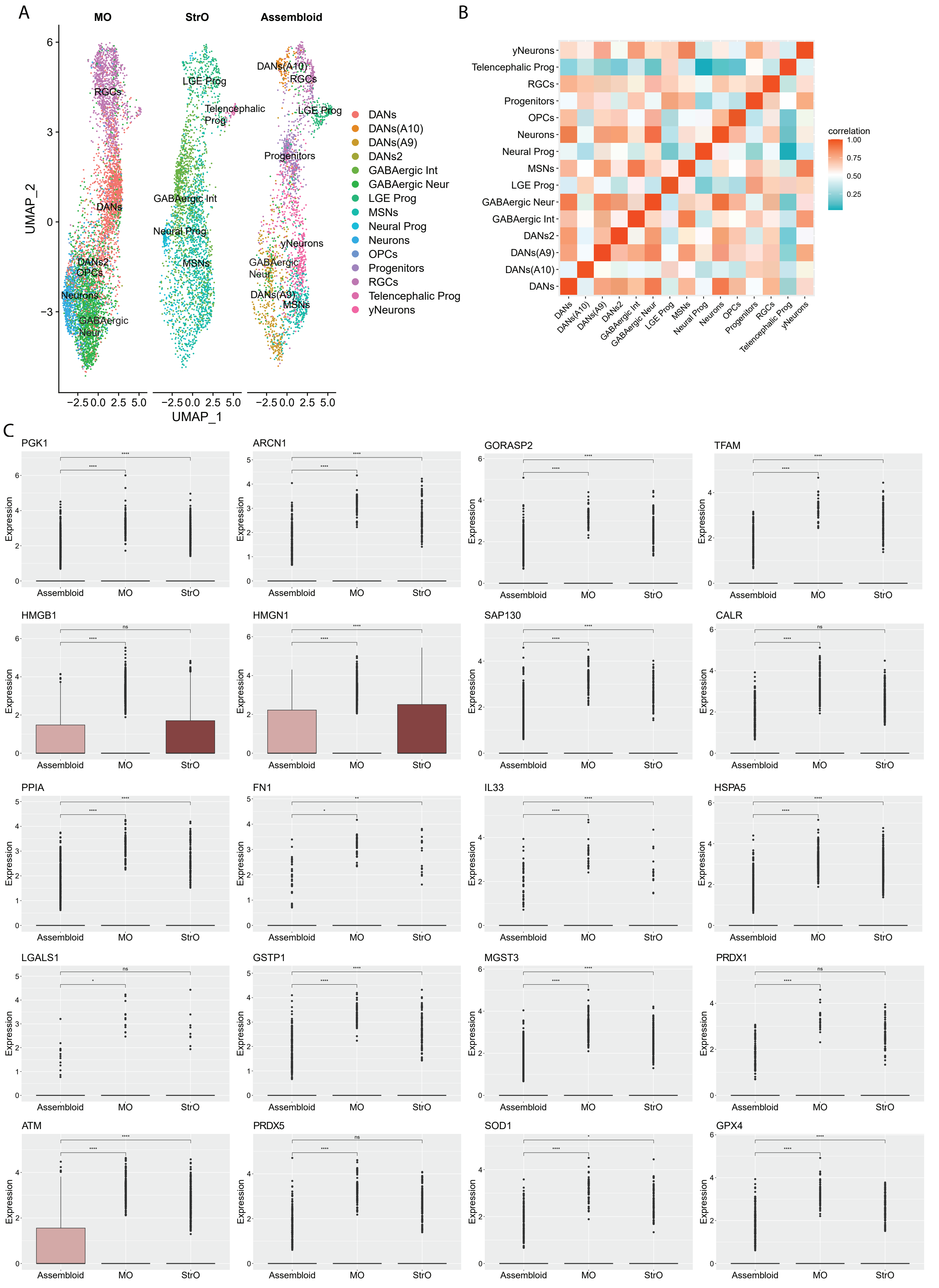


Supplementary Figure 10

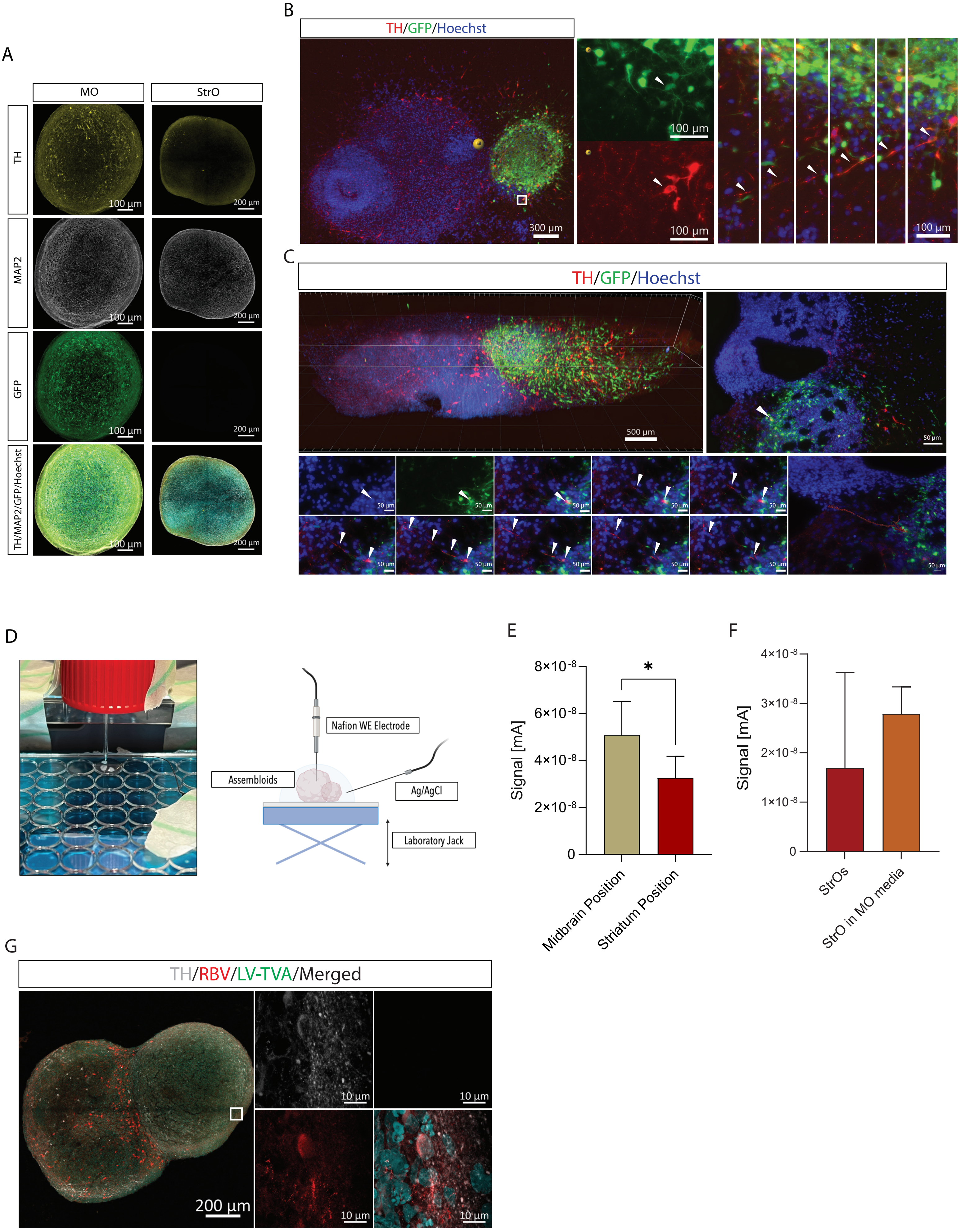


Supplementary Figure 11

A

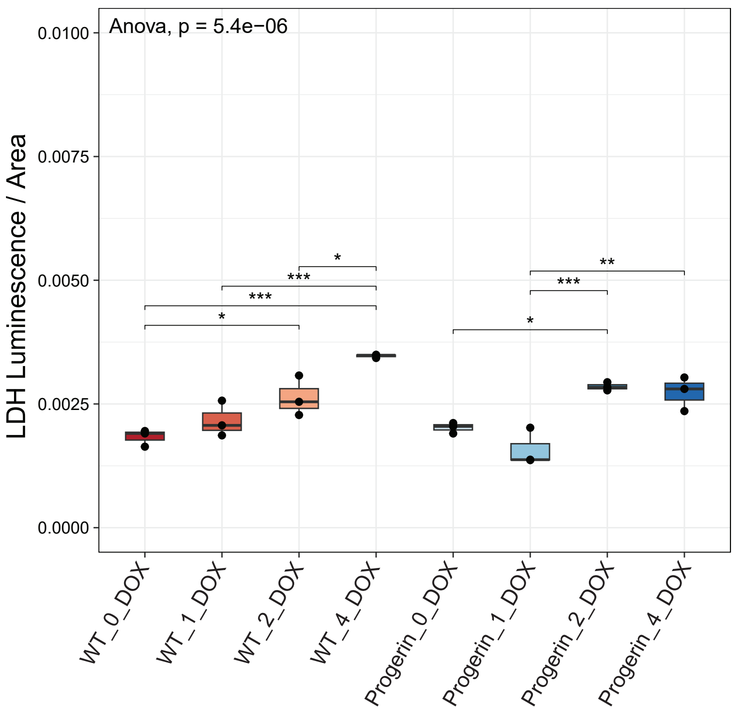


B

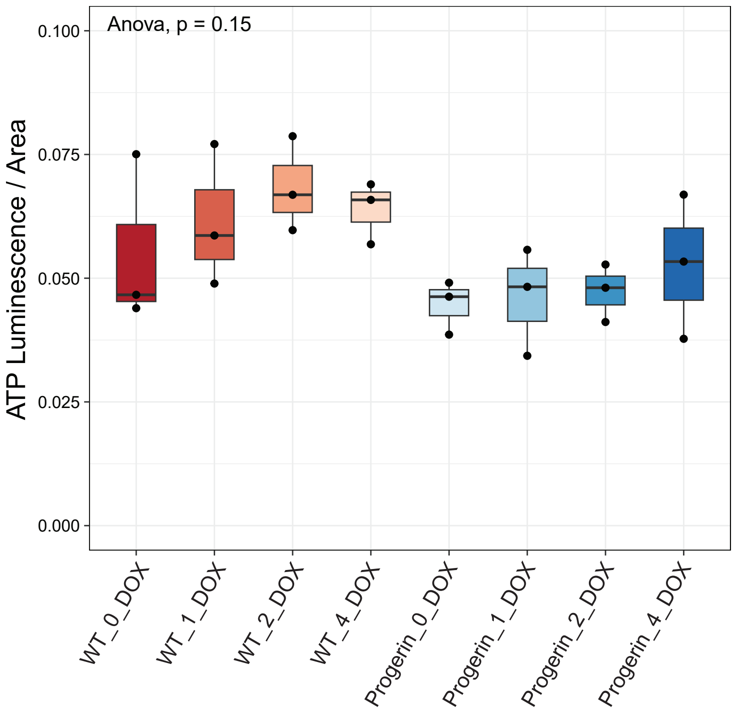


Supplementary Figure 12

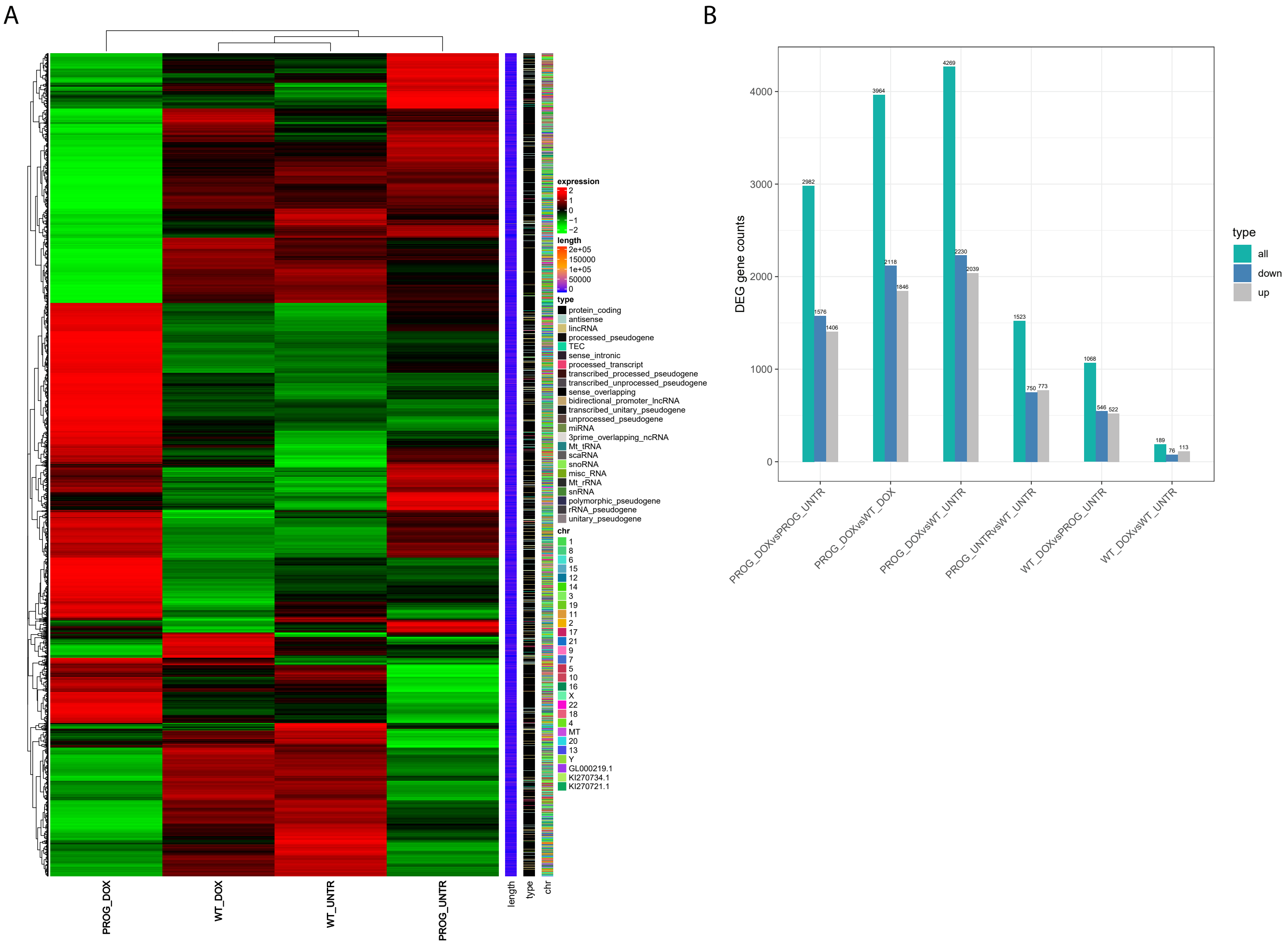


Supplementary Figure 13

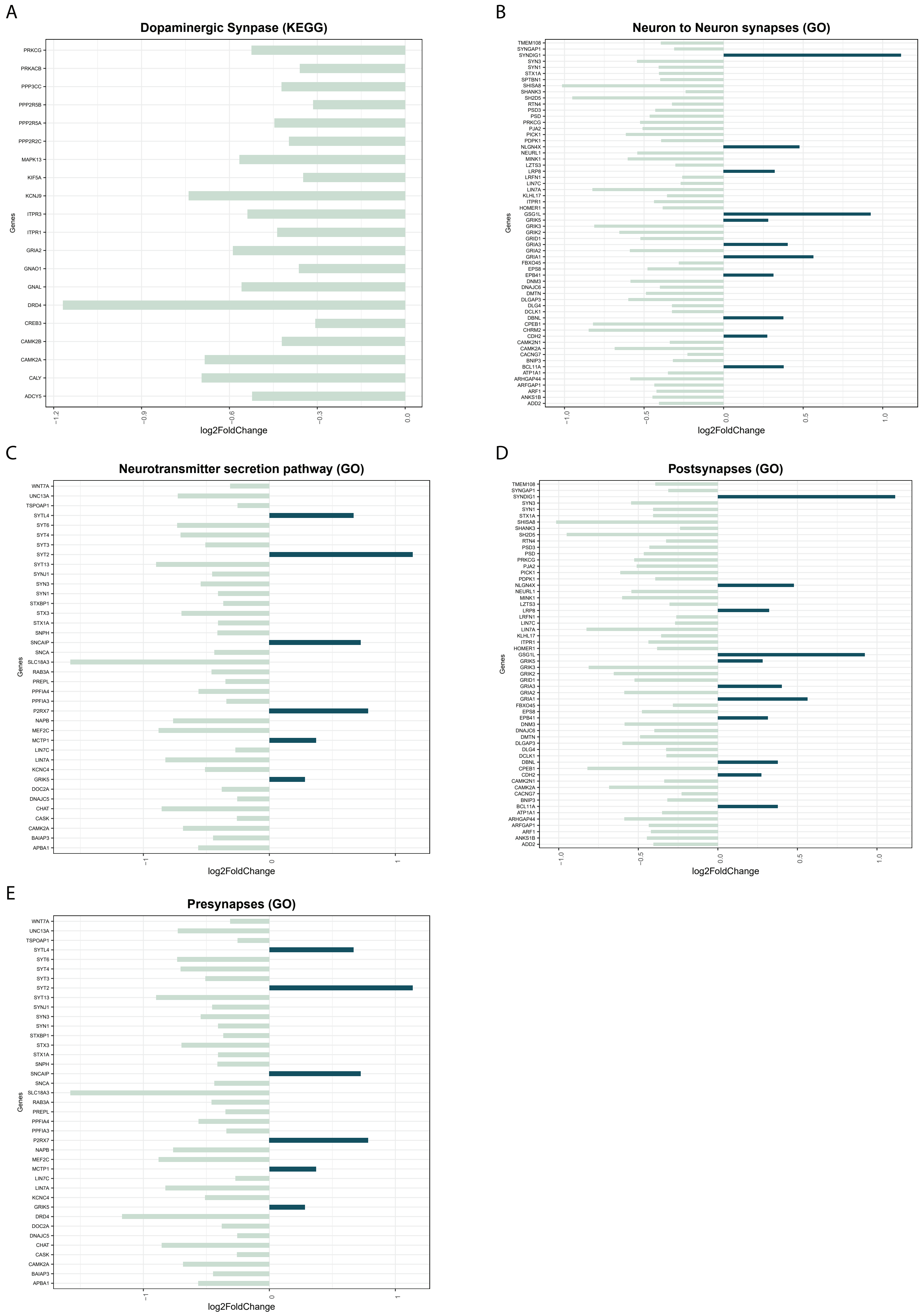


Supplementary Figure 14

**Supplementary Table 1: Cell lines used in the study.**

| **Lab identifier** | **Name** | **Sex** | **Age of sampling** | **Source** | **Reference Name** | **Karyotype** |
|---|---|---|---|---|---|---|
| 201 | WT 1 | F | - | GIBCO | A13777 | Normal |
| 200 | WT 2 | F | 83 | Reinhardt *et al.*, 2013 | K7.1 | Chr1-mosaic gain q21.2<br>Chr1- gain q25.3 |
| 389 | WT | M | 6 | Promocell | NHDF | Chr20q - gain |
| 390 | Progerin | M | 6 | Innsbruck, Prof. Frank Edenhover | NHDF progerin | Chr20q - gain |

**Supplementary Table 2: Primary and secondary antibodies used in Western blots.**

| Antibody | Source | Ref.no. | RRID | Species | Dilution |
|---|---|---|---|---|---|
| β-Actin | Cell Signaling | 3700S | *AB_2242334* | Mouse | 1:20000 |
| LAMINB1 | Abcam | ab16048 | *AB_443298* | Rabbit | 1:500 |
| LMNA | Sigma | L1293 | *AB_532254* | Rabbit | 1:500 |
| DRD1 | Abcam | ab216644 | *AB_2941932* | Rabbit | 1:300 |
| DRD2 | Abcam | ab85367 | *AB_10674739* | Rabbit | 1:300 |
| Histone H3 | Millipore | 05-1341 | *AB_1977240* | Mouse | 1:20000 |
| TUJ1 | BioLegend | 801201 | *AB_2313773* | Mouse | 1:20000 |
| TAU | Abcam | ab80579 | *AB_1603723* | Mouse | 1:1000 |
| Synaptotagmin1 | Synaptic Systems | 105011 | *AB_2619761* | Mouse | 1:200 |
| VAMP2 | Abcam | ab215721 | *AB_2923382* | Rabbit | 1:1000 |
| TH | Abcam | ab112 | *AB_297840* | Rabbit | 1:600 |
| DARPP32 | Abcam | ab40801 | *AB_731843* | Rabbit | 1:400 |
| GAD65 | R&D systems | AF2247 | *AB_2108039* | Goat | 1:1000 |
| ECL anti-goat | Santa Cruz | sc-2020 | *AB_631728* | Donkey | 1:1000 |
| Anti-rabbit H+L 800 | Cell Signaling | 5151 | *AB_10697505* | Goat | 1:10000 |
| Anti-mouse H+L 680 | Cell Signaling | 5470 | *AB_10696895* | Goat | 1:10000 |

**Supplementary Table 3: Primary and secondary antibodies used in immunofluorescence stainings.**

| Antibody | Source | Cat.no. | RRID | Species | Dilution |
|---|---|---|---|---|---|
| NESTIN | BD Bioscience | 611659 | *AB_399177* | Mouse | 1:600 |
| PAX6 | Biolegend | 901302 | *AB_2749901* | Rabbit | 1:300 |
| FOXA2 | Santa Cruz | sc-101060 | *AB_1124660* | Mouse | 1:100 |
| MASH1/ASCL1 | BD Bioscience | 556604 | *AB_396479* | Mouse | 1:200 |
| CORIN | R&D systems | MAB2209 | *AB_2082224* | Rat | 1:200 |
| TRA-1-60 | Millipore | MAB4360 | *AB_2119183* | Mouse | 1:50 |
| SSEA-4 | Millipore | MAB4304 | *AB_177629* | Mouse | 1:50 |
| NANOG | Millipore | AB5731 | *AB_2267042* | Rabbit | 1:200 |
| OCT4 | Abcam | ab19857 | *AB_445175* | Rabbit | 1:400 |
| CTIP2 | Abcam | ab18465 | *AB_2064130* | Rat | 1:300 |
| TH | Abcam | ab112 | *AB_297840* | Rabbit | 1:600 |
| MAP2 | Abcam | ab92434 | *AB_2138147* | Chicken | 1:1000 |
| SOX1 | R&D systems | AF3369 | *AB_2239879* | Goat | 1:100 |
| SOX2 | Abcam | ab97959 | *AB_2341193* | Rabbit | 1:200 |
| SOX2 | R&D Systems | AF2018 | *AB_355110* | Goat | 1:200 |
| DARPP32 | Abcam | ab40801 | *AB_731843* | Rabbit | 1:400 |
| P21 Waf1/Cip1 | Cell Signaling | 2946 | *AB_2260325* | Mouse | 1:200 |
| P16INK4a | Abcam | ab108349 | *AB_10858268* | Rabbit | 1:200 |
| P53 | Thermo Fisher Scientific | MA5-12557 | *AB_10989883* | Mouse | 1:200 |

| | | | | | |
|---|---|---|---|---|---|
| H2AX | Millipore | 05-636-I | *AB_2755003* | Mouse | 1:200 |
| 53BP1 | Novus | NB100-304 | *AB_10003037* | Rabbit | 1:200 |
| Anti-mouse 488 | Invitrogen | A21202 | *AB_141607* | Donkey | 1:1000 |
| Anti-mouse 568 | Invitrogen | A10037 | *AB_2534013* | Donkey | 1:1000 |
| Anti-goat 657 | Invitrogen | A21447 | *AB_2535864* | Donkey | 1:1000 |
| Anti-rabbit 647 | Invitrogen | A31573 | *AB_2536183* | Donkey | 1:1000 |
| Anti-chicken 568 | Invitrogen | A11041 | *AB_2534098* | Goat | 1:1000 |
| Anti-chicken 647 | Invitrogen | A21449 | *AB_2535866* | Goat | 1:1000 |
| Anti-mouse 488 | Invitrogen | A11029 | *AB_2534088* | Goat | 1:1000 |
| Anti-mouse 568 | Invitrogen | A11031 | *AB_144696* | Goat | 1:1000 |
| Anti-mouse 647 | Invitrogen | A21236 | *AB_2535805* | Goat | 1:1000 |
| Anti-rabbit 568 | Invitrogen | A11036 | *AB_10563566* | Goat | 1:1000 |
| Anti-rabbit 647 | Invitrogen | A21244 | *AB_2535812* | Goat | 1:1000 |

**Supplementary Table 4: Primers used in qPCRs for the striatum organoids characterization.**

| Gene | Forward Primer | Reverse Primer |
|---|---|---|
| DARPP32 | CCTGAAGGTCATCAGGCAGT | GGTCTTCCACTTGGTCCTCA |
| DRD1 | AGGGACATGTCTTTGGCTTCAG | GGGAACAGTGTTAGCACCTGTT |
| DRD2 | CTGAGGGCTCCACTAAAGGAG | CATTCTTCTCTGGTTTGGCG |
| CTIP2 | ATCCTCAGCCCCTTTTGTTT | GCCGTTGTTCCTGAATTGTT |
| ASCL1 | GTCCTGTCGCCCACCATCTC | CCCTCCCAACGCCACTGAC |
| FOXG1 | TGTTGACTCAGAACTCGCTGG | CTGCTCTGCGAAGTCATTGAC |
| FOXP1 | GCAGTTACAGCAGCAGCACCTCC | CAGCCTGGCCACTTGCATACACC |
| FOXP2 | AATGTGGGAGCCATACGAAG | GCCTGCCTTATGAGAGTTGC |
| OTX2 | TCAACTTGCCCGAGTCGAGG | CAATGGTCGGGACTGAGGTG |
| NKX1.2 | CGCATCCAATCTCAAGGAAT | TGTGCCCAGAGTGAAGTTTG |
| GSX2 | ATGTCGCGCTCCTTCTATGTC | ATGCCAAGCGGGATGAAGAAA |
| ACTINB | TCAAGATCATTGCTCCTCCTGAG | ACATCTGCTGGAAGGTGGACA |

**Supplementary Table 5:** Genes related to cellular and oxidative stress.

| Genes | Role in Cellular/Oxidative stress | References |
|---|---|---|
| PGK1 (Phosphoglycerate Kinase 1) | A protein kinase responsible for governing mitochondrial activity and cell stress-triggered autophagy. Downregulation has been linked to oxidative stress inhibition. | (Bhaduri et al., 2020; Xu et al., 2022) |
| ARCN1 (Archain 1) | Involvement in ER stress response. | (Bhaduri et al., 2020; Izumi et al., 2016) |
| GORASP2 (Golgi Reassembly Stacking Protein 2) | Involvement in ER stress response. | (Bhaduri et al., 2020; Zhang & Wang, 2018) |
| TFAM (Transcription Factor A, Mitochondrial) | Stress response induced by mitochondria dysfunction. | (Hunt et al., 2019) |
| HMGB1 (High Mobility Group Box 1) | Cytokine with a role in initiating neuroinflammation. | (Paudel et al., 2020) |
| HMGN1 (High Mobility Group Nucleosome Binding Domain 1) | Chromatin architectural protein. Can contribute to neuroinflammation. | (Farley et al., 2022; Furusawa & Cherukuri, 2010) |
| SAP130 (Sin3A Associated Protein 130) | It is increased in neuroinflammation. | (Y. Wang et al., 2019) |
| CALR (Calreticulin) | ER Ca2+ binding protein. Is activated by environmental stress. | (Chen et al., 2023; Michalak, 2023) |
| PPIA (Peptidylprolyl Isomerase A) | It has a role as molecular chaperone. Increased levels in ALS. | (Lauranzano et al., 2015) |
| FN1 (Fibronectin 1) | Extracellular matrix glycoprotein. Increased expression in cellular stress response. | (Dhanani et al., 2017) |
| IL33 (Interleukin 33) | Cytokine with a role in neuroinflammation. | (Sun et al., 2021; Zharichenko & Njoku, 2020) |
| HSPA5 (Heat Shock Protein Family A (Hsp70) Member 5) | Upregulated under ER stress for the clearing of misfolded proteins. | (Nowakowska et al., 2020) |
| LGALS1 (Galectin 1) | Highly expressed during neuroinflammation. | (Aalinkeel & Mahajan, 2016; J. Wang et al., 2015) |
| GSTP1 (Glutathione S−Tranferase) | Increased levels have been associated with cancer and neurodegenerative diseases. | (Allocati et al., 2018) |
| MGST3 (microsomal Glutathione S−Tranferase) | Possible role in neuroinflammation, with the production of LCT4 for the biosynthesis of proinflammatory LTs. | (Fetissov et al., 2002) |
| PRDX1 (Peroxiredoxin 1) | An antioxidant enzyme that protects the brain against oxidative stress. Increased levels have been found in AD patients. | (Kim et al., 2022; Szeliga, 2020) |
| ATM (ATM Serine/Threonine Kinase) | It is activated by oxidative stress and is important for the repair of DNA damage. | (Berger et al., 2017; Kozlov et al., 2016) |
| PRDX5 (Peroxiredoxin 5) | Antioxidant enzyme, protecting cells against oxidative stress (similar to PRDX1). | (Tavleeva et al., 2022; Yuan et al., 2004) |
| SOD1 (Superoxide Dismutase 1) | It is oxidative stress dependent and is upregulated during brain inflammation. | (Dell'Orco et al., 2021; Dimayuga et al., 2007) |
| GPX4 (Glutathione Peroxidase 4) | It is overexpressed during ferroptosis and neuroinflammation. | (Fang et al., 2023) |
